# Matrix game between full siblings in Mendelian populations

**DOI:** 10.1101/2024.03.04.583267

**Authors:** József Garay, Tamás Varga, Villő Csiszár, Tamás F. Móri, András Szilágyi

## Abstract

We demonstrate that static evolutionary stability implies the stability of the corresponding interior equilibrium point in genotype dynamics, while a certain form of monotonicity ensures the global stability of a homozygote state.

We apply our findings to familial selection in a diploid, panmictic population, where the survival rates of siblings within monogamous and exogamous families are determined by a matrix game, and the behavior is uniquely determined by an autosomal recessive-dominant or intermediate allele pair. We provide conditions for the existence of each homozygote.

In our numerical investigations of the Prisoner’s Dilemma between siblings, we distinguish two scenarios: cooperation (collaborating case) or defector-cooperator strategy pair (alternating case) that maximizes the siblings’ survival rates. Based on the stability of the pure cooperator and defector states, we provide a potential classification of genotype dynamics. We find that the pure cooperator population cannot fixate in the alternating case. However, in the collaborating case, fixation is possible but not necessary, since bistability, coexistence, moreover, the monostable fixation of pure defector state can also occur due to the interplay between the phenotypic payoff function and the genotype-phenotype mapping, which collectively determine the outcome of natural selection. In donation game, the classical Hamilton’s rule implies the fixation of the cooperation in all considered genotype-phenotype mappings.

**Author Summary:** In this article, we explore the dynamics among full siblings who can mutually aid each other for survival. The strategy (cooperator or defector) of each sibling is determined by Mendelian (dominant-recessive or intermediate) inheritance. The interaction between two siblings is modelled as a Prisoner’s Dilemma. We examine two scenarios: cooperation (collaborating case) and a defector-cooperator strategy pair (alternating case), aiming to maximize the combined survival rates of the interacting siblings. The endpoint of natural selection is determined by Mendelian inheritance and the two Prisoner’s dilemma scenarios. Our findings reveal the potential for the fixation of both cooperation and defection, as well as the stable coexistence of these strategies within the studied selection environment. Notably, in the alternating case, the fixation of a pure cooperating population is not achievable under the considered inheritance systems. Furthermore, when cooperation is recessive, its fixation is more likely but not guaranteed.

## Introduction

Our motivation is rooted in Haldane’s “*familial selection*” regime [1], where the size of a family is strictly limited by the available food resulting in the production of more newborns than can survive. This leads to intense struggle among the members of the same family [2]. In our study, we examine a slightly different scenario characterized by a frequency-dependent, survival matrix game between siblings, within monogamous and exogamous families [3-4].

In the context of sexual reproduction, organisms can be classified as either haploid (e.g., mosses) or diploid (e.g., mammals), depending on whether adult individuals are haploid or diploid. For our analysis, we focus on diploid organisms. Because haploid and diploid life cycles alternate during sexual reproduction, there are two approaches to modelling the evolution in diploid sexual populations: tracking either the haploid stage (gamete types) or the diploid stage (genotypes) across generations.

### Gene-centred models

These models typically adopt the gene pool approach, where the diploid parental population is assumed to be at Hardy–Weinberg equilibrium, and the focus is on the distribution of alleles [5-7].

### Genotype-centred models

When internal fertilization occurs and the survival rate of the juveniles depends on the mating pair, a genotype-centered population genetic model becomes necessary. The variable of these models is the distribution of diploid genotypes [3,8-15]. Although diploid embryos follow Hardy-Weinberg proportions under panmixia, the family-dependent survival selection can significantly alter the genotype proportions in adults. Consequently, the Hardy-Weinberg equilibrium is no longer useful for model formulation or analysis in these cases [7].

We mention two additional advantages of the genotype-centred models. Firstly, the classical Darwinian fitness is defined as the average growth rate of the phenotype. Consequently, the genotype centred model is the closest to the classical Darwinian view, since the genotype determines the phenotype. Secondly, kin selection theory focuses on the diploid individuals, the cost and benefit from their interactions, and the genetic relatedness between them [16]. In this sense, the classical Hamilton’s rule and the notion of inclusive fitness are also genotype-centred.

Genotype centred population genetic models have been used extensively to investigate the evolution of altruism within families [3,4, 8, 12, 14-15, 17]. We recall some key findings from these population genetic models. Firstly, the fixation of the altruistic behavior depends on the genotype-phenotype mapping [3, 14, 18]. Secondly, in the presence of additive cost and benefit functions, population genetic models yield the same results as Hamilton’s method [3-4, 16]. However, this equivalence does not hold in non-additive situations [4, 8]. These published models predominantly focus on the fixation of the altruistic homozygote. In contrast, our focus is on the evolutionary stability of a polymorphic diploid population, specifically examining the coexistence of different genotypes. Thus, we address the following two problems: 1. What are the static conditions for the existence of a stable mixed genotype distribution? 2. Do these conditions imply the dynamical stability of the corresponding genotype dynamics [3]?

Furthermore, altruism can be viewed as a one-person game, where an actor has two pure strategies (altruistic or selfish) and the recipient has no strategy. This prompts the question: Does the classical Hamilton’s rule hold true if the recipient also has a strategy, i.e., when the interaction between siblings is described by a two-person matrix game? As an application, we examine the case of two alleles at an autosomal locus, which uniquely determine the phenotypes under dominant-recessive or intermediate inheritance. We provide numerical examples where the Prisoner’s dilemma [19] and a particular version of it, the donation game [20], describe the interaction between the siblings.

## Results

### Genotype dynamics in sexual diploid population

Let us consider a very large, diploid population of *N* individuals. Here “very large” means that during sexual reproduction all possible genotypes manifest in the sexual population (with probability almost 1). The females and males differ only in their sex, with half-half sex ratio, and no sexual selection. We follow the change of the genotype frequencies in time. We consider *m* genotypes, so the distribution of genotypes belongs to the *m*-dimensional simplex

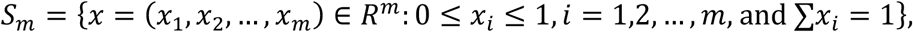

Our dynamics is based on the following assumptions.

#### Assumption A)

*The mating system is fixed* Let the probability of mating of genotypes *i* and *j* be *h*_*ij*_(*x*) ∈ [0,1], e.g. *h*_*ij*_(*x*) = *x*_*i*_*x*_*j*_ under panmixia.

#### Assumption B) The genotypes of the parents determine the genotypes of their offspring.

Denote by *p*_*k*(*ij*)_ ∈ [0,1] the probability that a given offspring of a mating pair of genotypes *i* and *j* has genotype *k*. Traditionally, all *p*_*k*(*ij*)_-s are collected in a mating table.

#### Remark 1.

Note that arbitrary genetic architecture can be considered here. For instance, the probability *p*_*k*(*ij*)_ can be calculated in the case of multi autosomal loci located on the same or different chromosomes, with or without recombination. Furthermore, if each allele can mutate to any other allele at the same locus, then each entry of the mating table is strictly positive.

Natural selection operates on phenotype level, thus we need proper information on the genotype-phenotype map.

#### Assumption C)

*The phenotype is genetically determined* Under this assumption, we can consider different selection regimes and calculate the survival rate of each sibling based on game-theoretical conflicts (see Remark 2). This condition ensures that the choice of strategy is not an individual decision but is solely determined by the genotype.

#### Assumption D)

*The survival rate of each offspring depends on the genotypes of the parents*, as the parents’ genotypes determine the genotypes of their offspring, which in turn determine their phenotypes. With assumptions C and D in place, we can consider survival games between siblings, which are determined by the genotypes of the parents. Let *n*_*k*(*ij*)_ denote the number of surviving offspring of genotype *k* from a mating pair of genotypes *i* and *j*.

#### Remark 2

The parameter *n*_*k*(*ij*)_ is general in the sense that *n*_*k*(*ij*)_ can incorporate various selection situations. For instance, the offspring number may depend on the genotypes of the parents [5] and the genotype distribution. The probability of survival for an offspring with genotype *k* and parents of genotypes *i* and *j* can also be influenced by the genotype distribution. Therefore, if the number of offspring from pairs with genotypes *i* and *j* is denoted by *a*_(*ij*)_(*x*), and *ρ*_*k*(*ij*)_(*x*) ∈ [0,1] is the probability of survival for an offspring with genotype *k* and with parents of genotypes *i* and *j* in a population of state *x*, then *n*_*k*(*ij*)_(*x*) can be calculated as *ρk*(*ij*) (*x*)*pk*(*ij*) *a*(*ij*) (*x*).

In summary, to follow the change in genotype distribution over time, we need to know the relative frequency of the mating pairs of genotypes *i* and *j*, i.e. *h*_*ij*_(*x*), and, for each mating pair of genotypes *i* and *j*, the number of surviving offspring with genotype *k*, i.e. *n*_*k*(*ij*)_, considering any distribution *x* within the set of possible parental genotypes *S*_*m*_.

The total number of offspring of genotype *k* produced by the entire parental population in state *x* can be expressed as follows:

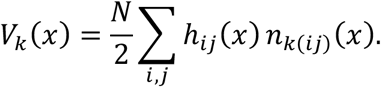

If *x* ∈ int*S*_*m*_, the *average production rate* of genotype *G*_*i*_ is *U*_*i*_(*x*) = *V*_*i*_(*x*)*⁄x*_*i*_, and the average production rate of the diploid population is *Ū* (*x*) = *∑*_*i*_ *V*_*i*_(*x*) = *∑*_*i*_ *x*_*i*_*U*_*i*_(*x*). To avoid degenerate situations, we make the assumption that *Ū* (*x*) > 0 for all *x*, that is, for any *x*, there exists at least one *i* such that *V*_*i*_(*x*) > 0.

## Genotype dynamics

Following the reasoning preceding the introduction of the replicator dynamics in Chapter 7.1 of Hofbauer and Sigmund [21], we consider a population that is sufficiently large and where generations blend continuously into each other. In such a scenario, it can be assumed that the frequency of a given phenotype is a differentiable function of time, so its change in time can be described by a system of differential equations. Analogously to the replicator dynamics, this change is expressed as the difference between the production rate of the given phenotype and the average production rate of the population:

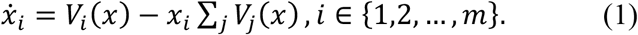

This dynamic process is referred to as *genotype dynamics* (for a detailed derivation, see SI C in [3]). The foundation of dynamics (1) lies in the basic tenet of Darwinism: the relative frequency of a given genotype increases when its production rate exceeds the average production rate of the entire population; in other words, when its *relative advantage over the whole population* is positive [22].

There is one key distinction between the replicator dynamics and the genotype dynamics: replicator dynamics applies to asexual populations, where parents of type *i* only produce offspring of the same type, whereas dynamics (1) models a sexual population, where parents of type *i* can produce offspring with genotypes other than *i* (see Tables 1, 4, 7 and 10).

**Table 1.** Genotype survival table based on the mating table for general matrix game. [a] and [A] are different alleles of the same gene. The phenotype of genotypes *G*_1_, *G*_2_ and *G*_3_ are in turn denoted by *s*_1_, *s*_2_ and *s*_3_. *x*_*i*_ is the frequency of the *G*_*i*_ genotype in the parental population. *p*_*k*(*ij*)_ is the probability that the genotype of a given offspring is *G*_*k*_ provided the parental genotypes are *G*_*i*_ and *G*_*j*_ , respectively. *n*_*k*(*ij*)_ is the number of offspring with genotype *G*_*k*_ provided the parental genotypes are *G*_*i*_ and *G*_*j*_ , respectively. “*A*” is a payoff matrix describing the interactions between siblings.

| Genotypes of parents | Average number of couples assuming panmixia | Average number of survived family members, for genotypes $G_1$, $G_2$, $G_3$ | | |
| --- | --- | --- | --- | --- |
| | | $G_1 = ([a],[a])$<br>$s_1$ | $G_2 = ([a],[A])$<br>$s_2$ | $G_3 = ([A],[A])$,<br>$s_3$ |
| $G_1 \times G_1$ | $\frac{N}{2} x_1^2$ | $p_{1(11)} = 1$<br>$n_{1(11)} = n s_1 A s_1$ | $p_{2(11)} = 0$<br>thus<br>$n_{2(11)} = 0$ | $p_{3(11)} = 0$<br>thus<br>$n_{3(11)} = 0$ |
| $G_1 \times G_2$ | $N x_1 x_2$ | $p_{1(12)} = \frac{1}{2}$<br>$n_{1(12)} = \frac{n}{4} (s_1 A s_1 + s_1 A s_2)$ | $p_{2(12)} = \frac{1}{2}$<br>$n_{2(12)} = \frac{n}{4} (s_2 A s_1 + s_2 A s_2)$ | $p_{3(12)} = 0$<br>thus<br>$n_{3(12)} = 0$ |
| $G_1 \times G_3$ | $N x_1 x_3$ | $p_{1(13)} = 0$<br>thus<br>$n_{1(13)} = 0$ | $p_{2(13)} = 1$<br>$n_{2(13)} = n s_2 A s_2$ | $p_{3(13)} = 0$<br>thus<br>$n_{3(13)} = 0$ |
| $G_2 \times G_2$ | $\frac{N}{2} x_2^2$ | $p_{1(22)} = \frac{1}{4}$<br>$n_{1(22)} = \frac{n}{16} (s_1 A s_1 + 2 s_1 A s_2 + s_1 A s_3)$ | $p_{2(22)} = \frac{1}{2}$<br>$n_{2(22)} = \frac{n}{8} (s_2 A s_1 + 2 s_2 A s_2 + s_2 A s_3)$ | $p_{3(22)} = \frac{1}{4}$<br>$n_{3(22)} = \frac{n}{16} (s_3 A s_1 + 2 s_3 A s_2 + s_3 A s_3)$ |
| $G_2 \times G_3$ | $N x_2 x_3$ | $p_{1(23)} = 0$<br>thus<br>$n_{1(23)} = 0$ | $p_{2(23)} = \frac{1}{2}$<br>$n_{2(23)} = \frac{n}{4} (s_2 A s_2 + s_2 A s_3)$ | $p_{3(23)} = \frac{1}{2}$<br>$n_{3(23)} = \frac{n}{4} (s_3 A s_2 + s_3 A s_3)$ |
| $G_3 \times G_3$ | $\frac{N}{2} x_3^2$ | $p_{1(33)} = 0$<br>thus<br>$n_{1(33)} = 0$ | $p_{1(33)} = 0$<br>thus<br>$n_{2(33)} = 0$ | $p_{1(33)} = 1$<br>$n_{3(33)} = n s_2 A s_2$ |

However, if the population is very large for the manifestation of all possible genotypes (i.e. *x*_*i*_ > 0 for every *i* = 1,2, … , *m*), then dynamics (1) can be rewritten in the following “replicator dynamics” form:

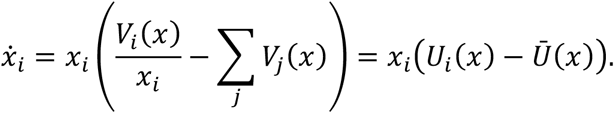

## Evolutionary stability of genotype distributions

Now we ask when a genotype distribution is an *evolutionarily stable genotype distribution* (ESGD) in an arbitrarily large sexual population. If every individual in a population has the same homozygous genotype, then following Maynard Smith and Price [23], we can consider this homozygous genotype to be evolutionarily stable if no mutant genotype can invade the population through natural selection. Mathematically, this means that the (relative) frequency of the resident homozygotes increases from generation to generation, provided mutation occurs rarely. However, the original concept of Maynard Smith and Price [23] is not directly applicable when there is polymorphism within the genotype population. Taking a dynamic perspective on stability, we consider a polymorphic state to be an ESGD if the system returns to this state after a small disturbance, such as a mutation [24]. In a static, evolutionary formulation, a genotype distribution is referred to as an ESGD if the average production rate of the entire genotype population (as seen on the right-hand side of inequality (2) below) is lower than that of the genotype subpopulation at ESGD (on the left-hand side of inequality (2)). In other words, the subpopulation in the ESGD state has a relative advantage over the entire population.

### Definition 1

*A genotype distribution x*^*^ ∈ int *S*_3_ *is evolutionary stable if*

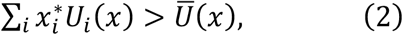

*provided that x is sufficiently close to x*^*^.

Clearly, inequality (2) can be rewritten as

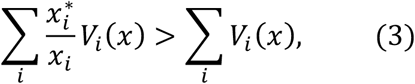

and we immediately see that *x*^*^ ∈ int*S*_3_ is an ESGD if and only if

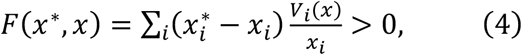

whenever *x* is close enough to *x*^*^. We note, here *V*_*i*_ (*x*)*⁄x*_*i*_ gives the average production rate of genotype *G*_*i*_. Thus *V*_*i*_ (*x*)*⁄x*_*i*_ is similar to the notion of fitness in asexual populations.

In asexual populations, the different coexisting types exhibit the same fitness in an evolutionarily stable state. In our setup, a similar claim holds. In SI A, we show that the following equilibrium condition (Nash equilibrium, in game theoretical terms) can be given.

## Equilibrium condition

If *x*^*^ ∈ int *S*_3_ is an ESGD and *x* is an arbitrary state in *S*_3_ then

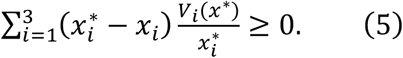

Furthermore, one of the characteristic properties of interior Nash equilibria in asexual matrix games holds true in this model as well. Namely, in interior Nash equilibria the average production rate of different genotypes must be equal. Formally, let *x*^*^ ∈ int*S*_*m*_ be an ESGD. Then, for every *i, j* ∈ {1,2, … , *m*} we have

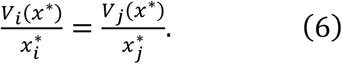

We note that equation (6) implies that an ESGD is a rest point of the genotype dynamics (1).

Following the argument in Chapter 7.2 of Hofbauer and Sigmund [21] we can prove the ensuing result (see Theorem SI.1 in SI B).

### Theorem 1

*Assume that x*^*^ ∈ int *S*_*m*_ *is an ESGD. Then x*^*^ *is a locally asymptotically stable rest point of genotype dynamics* (1).

We also need to define ESGD for a vertex of the simplex, particularly when every individual has the same homozygous genotype. This concept was introduced in SI C in [3], and we will now recall that definition. We say that *x*^*^ = (1,0, … ,0) is an ESGD if *V*_1_(*x*) > *x*_1_ *∑*_*j*_ *V*_*j*_ (*x*) for every *x* close enough to *x*^*^. It has been proven that if the vertex *x*^*^ = (1,0, … ,0) is an ESGD, then it is a locally asymptotically stable equilibrium point of the genotype dynamics (1) (see SI C in [3]).

## Application: Filial selection with matrix game between full siblings

Let us consider a diploid species with internal fertilization. Our assumptions are as follows: Mutations occur rarely enough. The species practices monogamy, resulting in a relatedness of 1/2 between full siblings. We consider *panmixia*, meaning that the probability of mating between genotypes *i* and *j* is *h*_*ij*_(*x*) = *x*_*i*_*x*_*j*_. The population size *N* is very large, with *N*/2 random couples formed, ensuring random mating and preventing inbreeding. As claimed in the introduction, our aim is to generalize Haldane’s concept of “*familial selection*” [1]. To achieve this, we also use the simplifying assumptions that each couple breeds a fixed number *n* of offspring, i.e. *a*_*ij*_(*x*) = *n* for all *i, j*. Furthermore, an offspring’s survival probability depends only on the behavior of its siblings and is independent of the overall genotype distribution in the population. These simplifications ensure that the frequency-dependent interaction between mating pairs and the population-dependent juvenile survival rate together cannot mask the effect of the survival game between full siblings. Due to the differing survival probabilities of distinct genotypes, the diploid parental population, in general, deviates from Hardy-Weinberg equilibrium.

For simplicity, we only consider one autosomal locus with two alleles, denoted by *a* and *A*, respectively, so three genotypes, *G*_1_ = ([*a*], [*a*]), *G*_2_ = ([*a*], [*A*]), and *G*_3_ = ([*A*], [*A*]) are possible. Their frequencies are denoted by *x*_1_, *x*_2_ and *x*_3_. Then the state of the population, represented by the vector *x* = (*x*_1_, *x*_2_, *x*_3_), belongs to

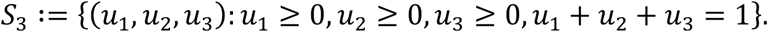

### Genotype-phenotype mapping

We assume that the genotypes uniquely determines the phenotypes. We denote by *s*_1_, *s*_2_ and *s*_3_ the phenotype of genotype *G*_1_ = ([*a*], [*a*]), *G*_2_ = ([*a*], [*A*]) and *G*_3_ = ([*A*], [*A*]), respectively. We will consider the dominant-recessive and intermediate Mendelian inheritance systems.

We assume that there is only game theoretical interaction between full siblings which is described by a one-shot matrix game and the entries of the payoff matrix are survival probabilities. This means that each sibling’s survival rate is the average individual payoff from its possible interactions with its siblings. Accordingly, we consider a two-dimensional matrix game within each family, with the same payoff matrix *A* ∈ *R*^2*×*2^,

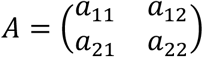

where *a*_*ij*_ ∈ (0,1] for every *i, j* ∈ {1, 2}.

The phenotypes are pure or mixed strategies from the simplex

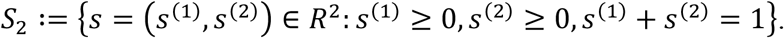

Under recessive–dominant inheritance, each individual adopts a pure strategy, either (1,0) or (0,1). Under intermediate inheritance, the homozygotes follows a pure strategy once again, while each heterozygote employs the mixed strategy (1*⁄*2 , 1*⁄*2). Note that the survival rate of offspring depends solely on their own phenotype and the phenotypes of their full siblings, which are determined by their parents’ genotypes. Consequently, the family unit, as a genetically well-defined interacting group, introduces a “group selection ingredient” to our model.

## Arbitrary inheritance

First, let us provide an overview of payoffs in the general case for two alleles at a single locus (see Table 1). Then, genotype *G*_1_ = ([*a*], [*a*]) has phenotype *s*_1_, genotype *G*_2_ = ([*a*], [*A*]) has phenotype *s*_2_ and genotype *G*_3_ = ([*A*], [*A*]) has phenotype *s*_3_. In other words, every heterozygote exhibits the same phenotype, regardless of the parent from which the allele are derived.

Above we have given the specification of a selection regime for Haldane’s familial selection model, when a two-person matrix game describes the interaction between full siblings. In this application, the number of surviving offspring *n*_*k*(*ij*)_ are listed in Table 1.

Within each family the game theoretical interaction is well mixed, meaning that the genotypes of the offspring are independent, and the pairs that interact are selected by chance. Therefore, the types of players in a given pair are also independent. For instance, in a *G*_2_*×G*_2_ family there are genotypes *G*_1,_ *G*_2_ and *G*_3_, with probabilities of 1/4, 1/2 and 1/4, respectively. As a result, a *G*_1_ juvenile interacts with phenotypes *s*_1_, *s*_2_ and *s*_3_ with probabilities of 1/4, 1/2 and 1/4, respectively. Consequently, the survival rate of a focal *G*_1_ juvenile is 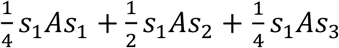 . Since the birth rate of a *G*_1_ juvenile in a *G*_2_*×G*_2_ family is 1/4, the average (expected) number of surviving *G*_1_ juveniles in *G*_2_*×G*_2_ families is 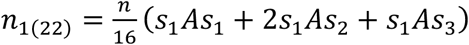. (For more details see SI C.)

Based on Table 1, the total number of individuals of genotypes *G*_1_, *G*_2_ and *G*_3_ in the next generation can be calculated as follows:

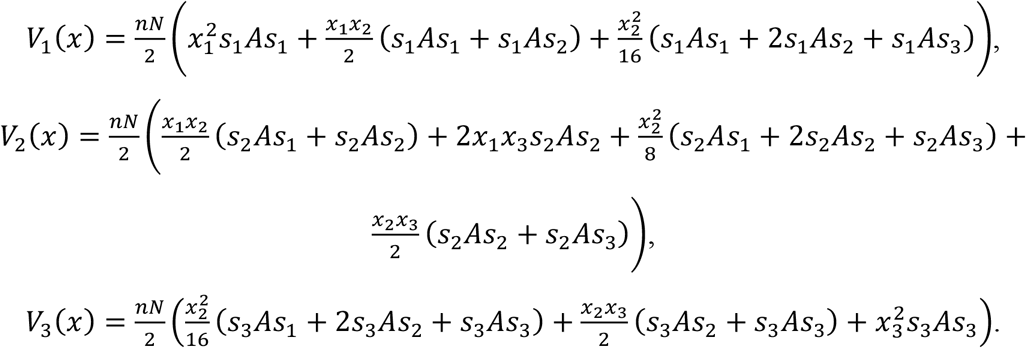

Using these production rates above, it is easy to get the actual version of the genotype dynamics (1) for the considered selection situation.

We focus on the following questions in the general case when the phenotype of *G*_1_ homozygotes is (1,0) while the phenotype of *G*_3_ homozygotes is (0,1).

**Question 1**. What is a necessary condition for the stability of the homozygote sates? (Here we only mention the first order condition. The detailed mathematical answer can be found in SI C.) Let allele [a] be recessive while allele [A] is dominant.

State (1,0,0), that is, where every individual is a recessive homozygote *G*_1_ = ([*a*], [*a*]), is a local ESGD if

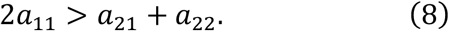

State (0,0,1), that is, where every individual is a dominant homozygote *G*_3_ = ([*A*], [*A*]), is a local ESGD if

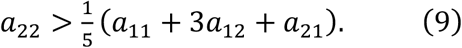

**Question 2**. What is a necessary condition for the coexistence of all the genotypes?

Clearly, the homozygote states are the only possible rest points of the genotype dynamics on the border of the simplex *S*_3_, and *S*_3_ is positively invariant respect to the genotype dynamics (see SI C in [3]). Thus, according to the two-dimensional Poincaré–Bendixson theorem, if these two rest points are repellors, then (in biological terms) we have either stable or cyclic coexistence of the two phenotypes.

State (1,0,0), that is, where every individual is a recessive homozygote *G*_1_, is a repellor (see SI C, Theorem SI.5), if

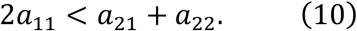

State (0,0,1), that is, where every individual is a dominant homozygote *G*_3_, is a repellor (see SI C, Theorem SI.6), if

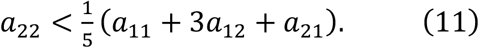

We note that the fulfillment of inequalities (10) and (11) imply that the homozygote states are not ESGD-s, thus based on our previous findings (see SI C in [3]), both states are repellor. Therefore, the combination of these two inequalities provides a sufficient condition for the coexistence of all genotypes (for additional details, see SI D). It is worth noting that conditions (8–11) are applicable to arbitrary payoff matrices, as long as phenotype *s*_1_ is recessive.

### Remark 3.

For the convenience of readers, we give a useful aid for the investigation of an arbitrary payoff matrix. Condition (2) of ESGD is nonlinear in the genotype distribution (SI C in [3]) in the neighborhood of the altruistic and selfish homozygote vertices. In the case where the linear term of the function *F*(*x*^*^, *x*) (see equation (4)) is not zero, its sign determines the evolutionary stability (positive linear term) or instability (negative linear term) of the homozygote vertex in question. When the linear term is zero, we have to check the second order terms of the function *F*(*x*^*^, *x*) and this investigation is not self-evident. The reader may think this case rather special thus less interesting. This is not so, since, as we have already observed, if the altruistic allele is dominant, the linear term is zero in the neighborhood of the altruistic homozygote vertex [3]. The reason that in the neighborhood of altruistic homozygote vertex the most frequent family is *G*_1_*×G*_1_, the family with the second largest frequency is *G*_1_*×G*_*2*_, and the other families are very rare, thus the production rates of different genotypes depend on what happens in these two families. But when the altruistic allele is dominant, in both families there are only altruistic individuals, thus the first order term vanishes. Similarly, when the selfish allele is dominant, the linear term vanishes in the neighborhood of the selfish homozygote vertex. We note that this situation is similar to that in the classical definition of ESS for matrix games [21], where the stability (i.e. second order) condition is needed for the evolutionary stability of a mixed Nash equilibrium.

Since the second order condition is often quadratic in terms of the entries of the payoff matrix, we also present simple linear inequalities implying that the vertex is (or at least may be) a repellor (see Tables 2, 3). This is the main reason why we introduced the genotype dynamics, since using differential equations the stability of the vertices can be investigated with success.

**Table 2.**
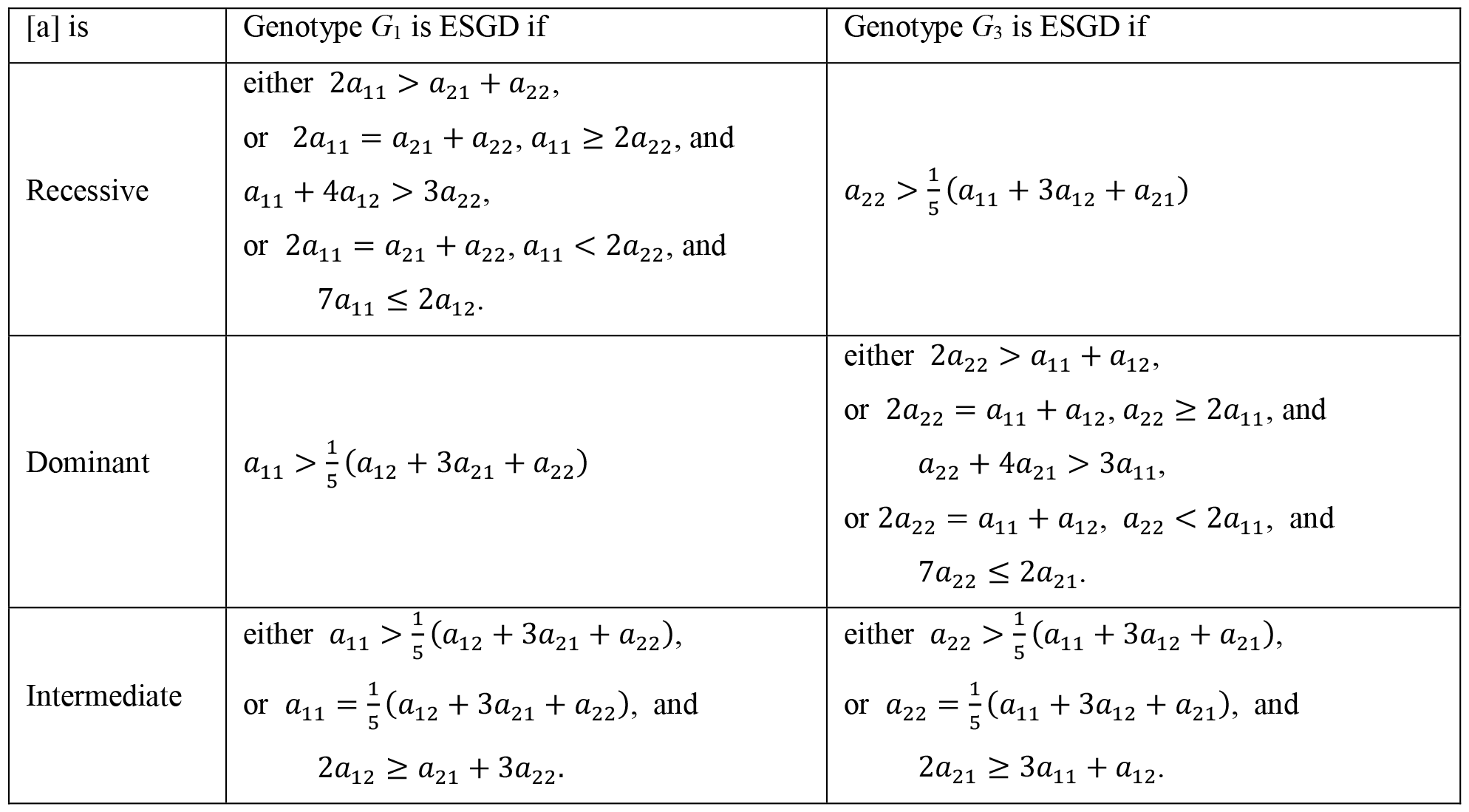
Conditions when states (1,0,0) and (0,0,1), respectively, are ESGD-s. (1,0,0) is the state when the genotype of every individual is *G*_1_=([a],[a]) with phenotype (1,0). (0,0,1) is the state when the genotype of every individual is *G*_3_=([A],[A]) with phenotype (0,1). [a] and [A] are different alleles of the same gene. (*a*_*ij*_)_2*×*2_ is a 2×2 payoff matrix describing the interactions between siblings.

| [a] is | Genotype $G_1$ is ESGD if | Genotype $G_3$ is ESGD if |
| --- | --- | --- |
| Recessive | either $2a_{11} > a_{21} + a_{22}$,<br>or $2a_{11} = a_{21} + a_{22}$, $a_{11} \geq 2a_{22}$, and<br>$a_{11} + 4a_{12} > 3a_{22}$,<br>or $2a_{11} = a_{21} + a_{22}$, $a_{11} < 2a_{22}$, and<br>$7a_{11} \leq 2a_{12}$. | $a_{22} > \frac{1}{5}(a_{11} + 3a_{12} + a_{21})$ |
| Dominant | $a_{11} > \frac{1}{5}(a_{12} + 3a_{21} + a_{22})$ | either $2a_{22} > a_{11} + a_{12}$,<br>or $2a_{22} = a_{11} + a_{12}$, $a_{22} \geq 2a_{11}$, and<br>$a_{22} + 4a_{21} > 3a_{11}$,<br>or $2a_{22} = a_{11} + a_{12}$, $a_{22} < 2a_{11}$, and<br>$7a_{22} \leq 2a_{21}$. |
| Intermediate | either $a_{11} > \frac{1}{5}(a_{12} + 3a_{21} + a_{22})$,<br>or $a_{11} = \frac{1}{5}(a_{12} + 3a_{21} + a_{22})$, and<br>$2a_{12} \geq a_{21} + 3a_{22}$. | either $a_{22} > \frac{1}{5}(a_{11} + 3a_{12} + a_{21})$,<br>or $a_{22} = \frac{1}{5}(a_{11} + 3a_{12} + a_{21})$, and<br>$2a_{21} \geq 3a_{11} + a_{12}$. |

**Table 3.**
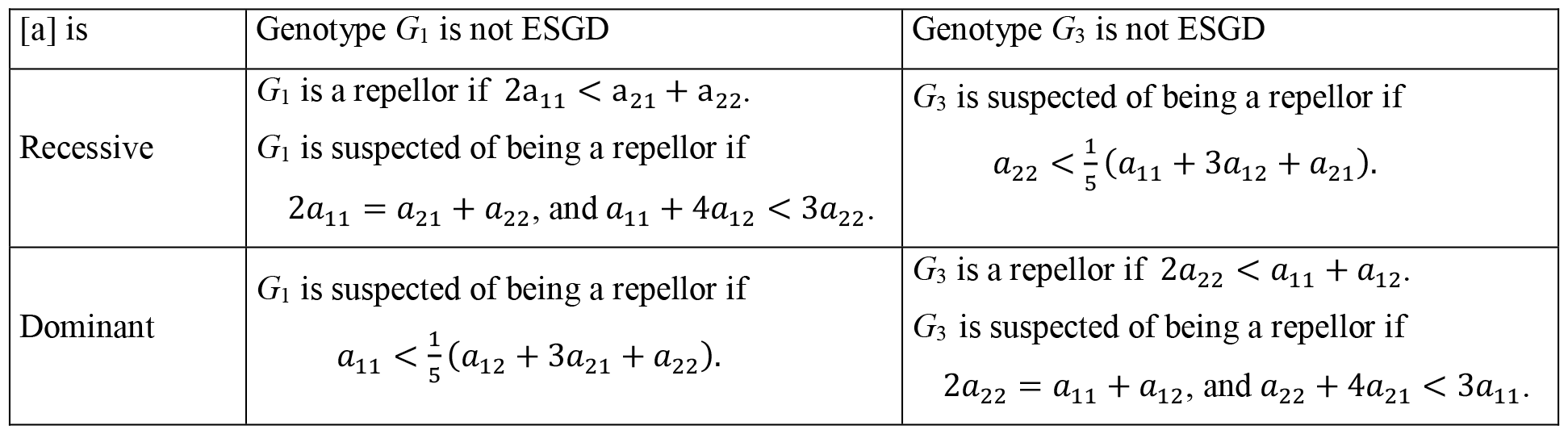
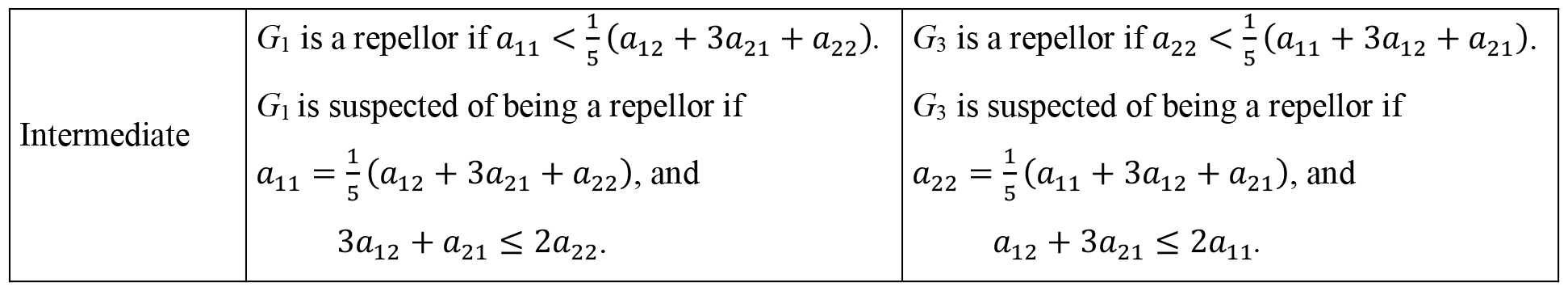
Conditions when states (1,0,0) and (0,0,1), respectively, are repellors. (1,0,0) is the state when the genotype of every individual is *G*_1_=([a],[a]) with phenotype (1,0). (0,0,1) is the state when the genotype of every individual is *G*_3_=([A],[A]) with phenotype (0,1). [a] and [A] are different alleles of the same gene. (*a*_*ij*_)_2*×*2_ is a 2×2 payoff matrix describing the interactions between siblings.

In the following, we aim at illustrating how the combination of the phenotypic payoff matrix and the genotype-phenotype mapping determines the solution of the genotype dynamics. It is important to note that our intention is not to provide a topological classification for the genotype dynamics, but rather to offer some numerical examples for better understanding. For this purpose, we assume that the game between siblings is a Prisoner’s dilemma, thus we consider cooperator and defector pure strategies and the following payoff matrix:

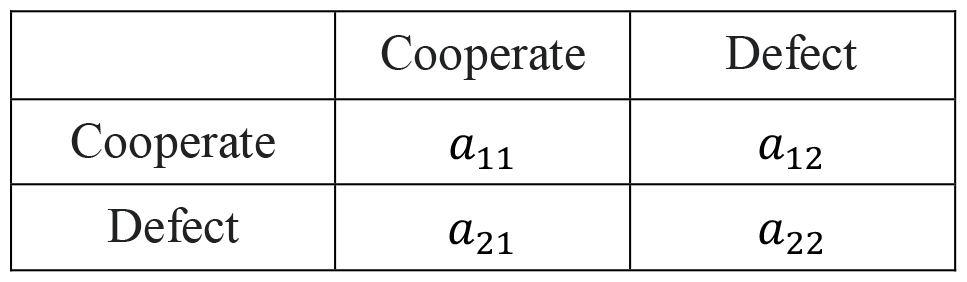

where *a*_21_ > *a*_11_ > *a*_22_ > *a*_12_. From the viewpoint of group selection, we distinguish two subclasses of prisoner’s dilemma games according to how siblings can maximize the “welfare of family” (i.e. the sum of their payoffs).

### Collaborating case

If 2*a*_11_ > *a*_21_ *+ a*_12_, then both siblings will cooperate in order to maximize their additive survival rate. Note that although the iterated prisoner’s dilemma game requires this condition, we concentrate only on the one-shot game.

### Alternating case

If 2*a*_11_ *< a*_21_ *+ a*_12_, then one of the siblings will cooperate while the other one will defect to maximize their respective additive survival rates.

We emphasize that in the above inequalities, the multiplier 2 is unrelated to the degree of relatedness between full siblings but only refers to the number of players involved.

In the following subsection we analyze the above familial selection models depending on whether the cooperator phenotype is recessive, dominant or intermediate.

## The cooperator phenotype is recessive

If the cooperator phenotype is recessive then *s*_1_ = (1,0) and *s*_2_ = *s*_3_ = (0,1). For the relating table, see Table 4.

**Table 4.**
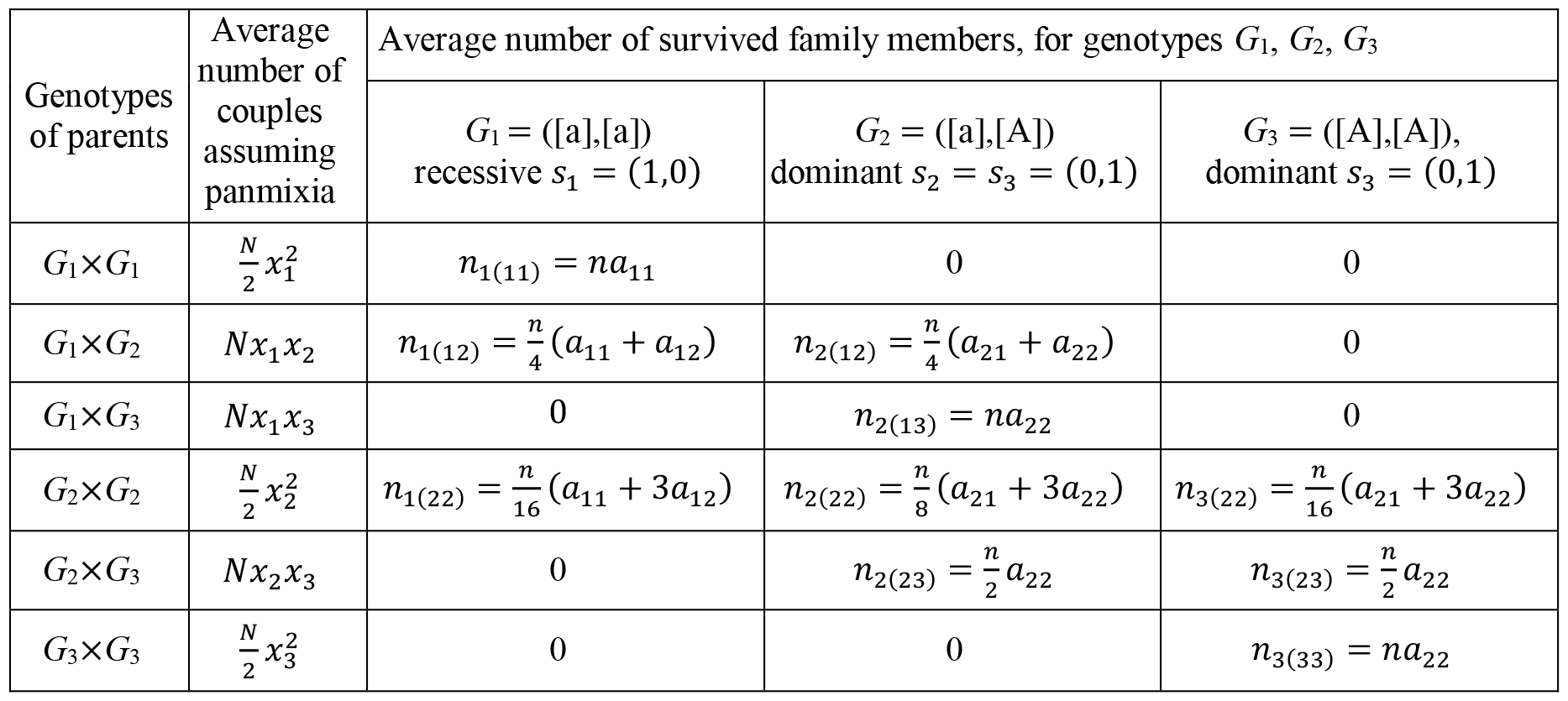
Genotype survival table based on Table 1 if the cooperator strategy is recessive. The phenotype of genotype *G*_1_ is the cooperator strategy represented by *s*_1_ = (1,0), the phenotype of genotype *G*_2_ and *G*_3_ is the defector strategy represented by *s*_3_ = (0,1). *s*_1_ is recessive while *s*_3_ is dominant. *p*_*k*(*ij*)_ is the probability that the genotype of a given offspring is *G*_*k*_ provided the parental genotypes are *G*_*i*_ and *G*_*j*_ , respectively. *n*_*k*(*ij*)_ is the number of offspring with genotype *G*_*k*_ provided the parental genotypes are *G*_*i*_ and *G*_*j*_ , respectively. “*A*” is a payoff matrix describing the interactions between siblings.

Here, inequalities (8–9) ensure the local stability of the homozygous states (1,0,0) and (0,0,1) while inequalities (10–11) provide sufficient conditions for coexistence. Based on this, we can identify four subcases in the collaborating case: both homozygous states are stable, one of them is stable, or both of them are unstable. In the alternating case, however, (1,0,0) can never be stable so only two subcases are possible: (1,0,0) is unstable and (0,0,1) is stable or both homozygous states are unstable. These subcases are illustrated with particular examples in Tables 5 and 6.

**Table 5.**
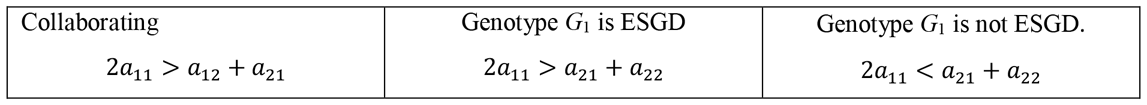
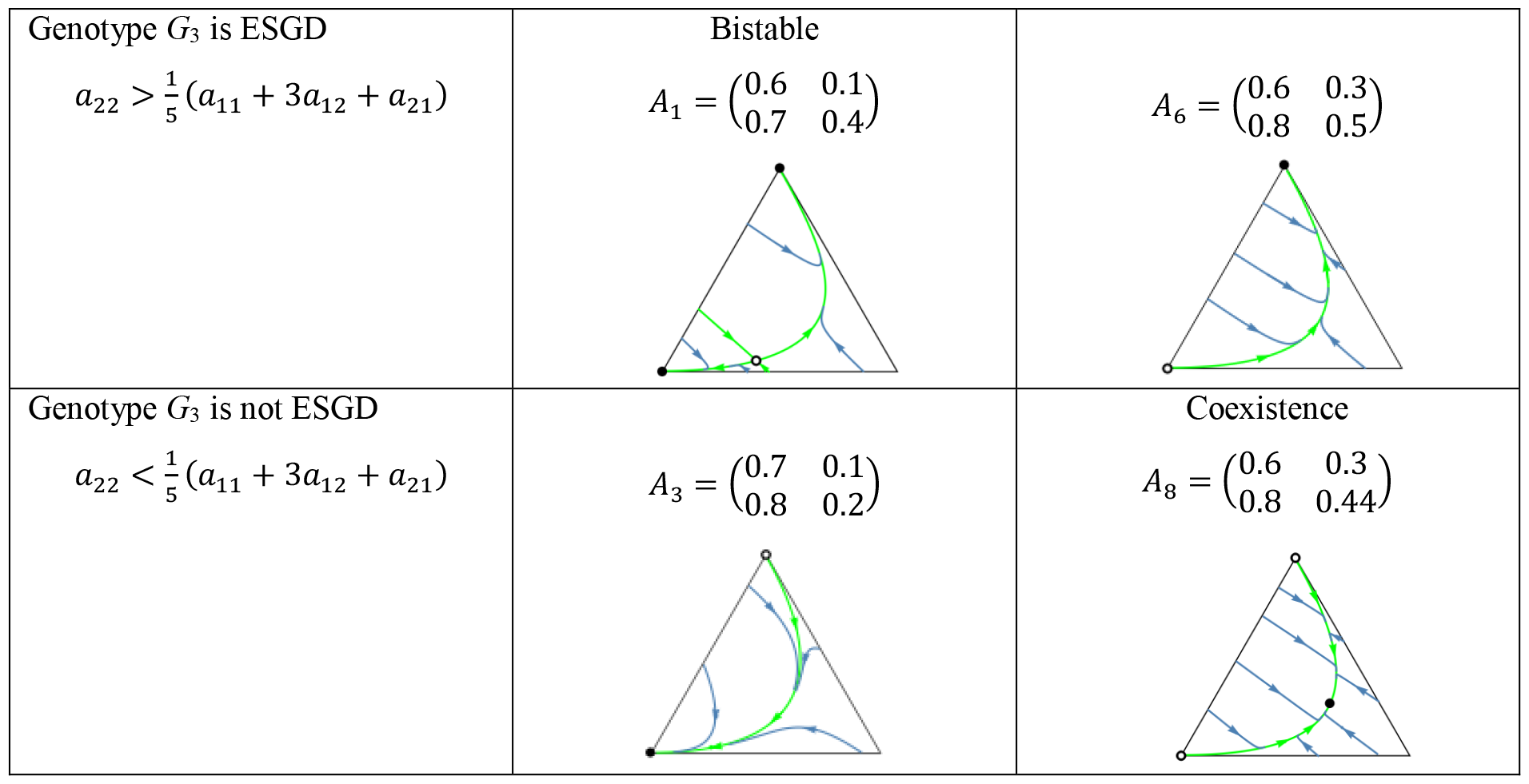
Examples where cooperation is recessive in collaborating PD game. Black dots correspond to an asymptotically stable rest points while empty dots represent unstable equilibrium points. The bottom-left vertex of the triangle corresponds to the (1,0,0) state, in which all individuals are ([a],[a]) cooperator homozygotes. The bottom-right vertex corresponds to the (0,1,0) state, where all individuals are ([a],[A]) heterozygotes. The top vertex corresponds to the (0,0,1) state, where all individuals are ([A],[A]) defector homozygotes. Black dots and circles denote asymptotically stable and unstable rest point, respectively.

| Collaborating | Genotype $G_1$ is ESGD | Genotype $G_1$ is not ESGD. |
| --- | --- | --- |
| $2a_{11} > a_{12} + a_{21}$ | $2a_{11} > a_{21} + a_{22}$ | $2a_{11} < a_{21} + a_{22}$ |
| Genotype $G_3$ is ESGD<br>$a_{22} > \frac{1}{5}(a_{11} + 3a_{12} + a_{21})$ | Bistable<br>$A_1 = \begin{pmatrix} 0.6 & 0.1 \\ 0.7 & 0.4 \end{pmatrix}$ | $A_6 = \begin{pmatrix} 0.6 & 0.3 \\ 0.8 & 0.5 \end{pmatrix}$ |
| Genotype $G_3$ is not ESGD<br>$a_{22} < \frac{1}{5}(a_{11} + 3a_{12} + a_{21})$ | $A_3 = \begin{pmatrix} 0.7 & 0.1 \\ 0.8 & 0.2 \end{pmatrix}$ | Coexistence<br>$A_8 = \begin{pmatrix} 0.6 & 0.3 \\ 0.8 & 0.44 \end{pmatrix}$ |

**Table 6.**
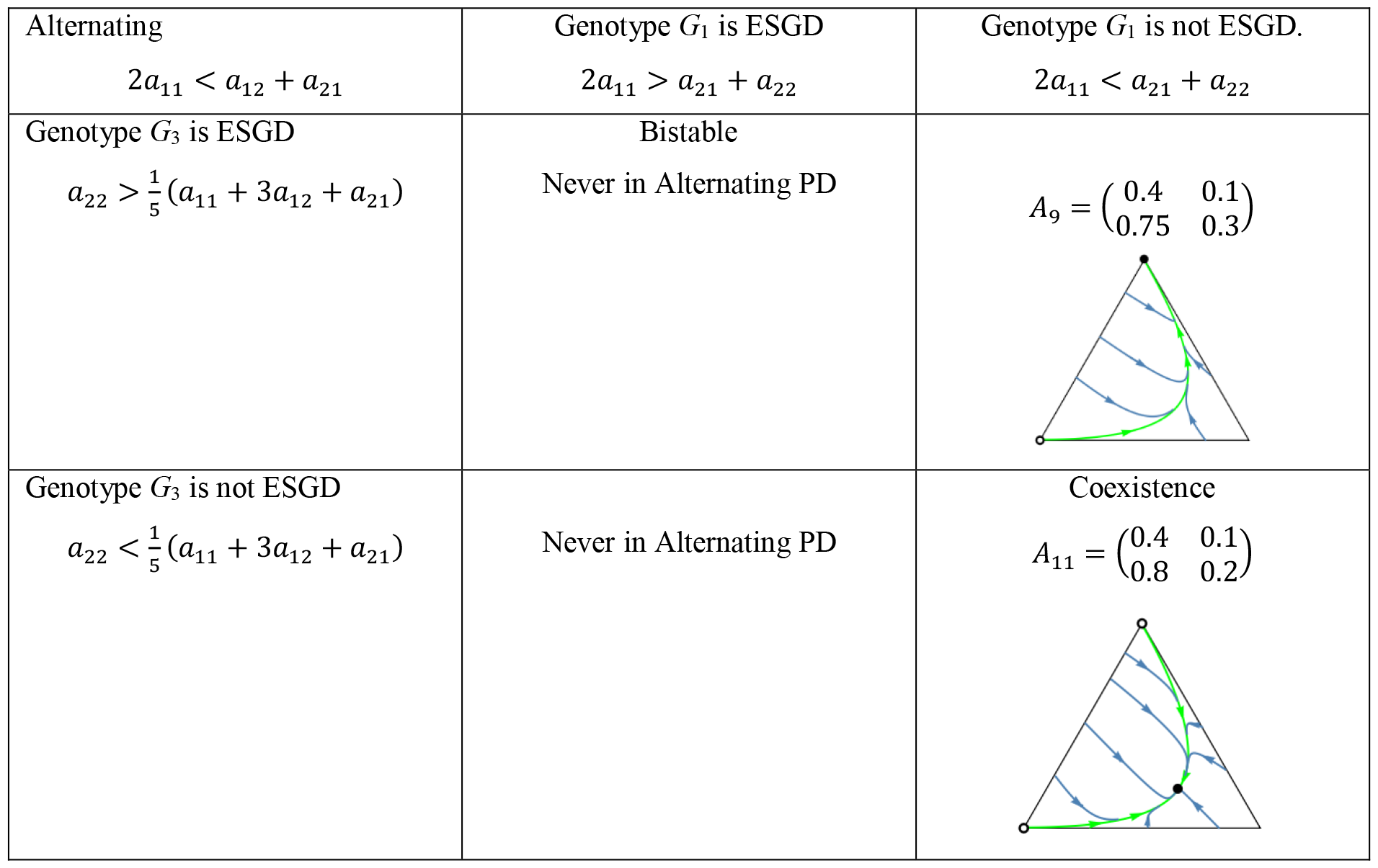
Examples where cooperation is recessive in alternating PD game. See the caption at Table 5.

| Alternating<br>$2a_{11} < a_{12} + a_{21}$ | Genotype $G_1$ is ESGD<br>$2a_{11} > a_{21} + a_{22}$ | Genotype $G_1$ is not ESGD.<br>$2a_{11} < a_{21} + a_{22}$ |
| --- | --- | --- |
| Genotype $G_3$ is ESGD<br>$a_{22} > \frac{1}{5}(a_{11} + 3a_{12} + a_{21})$ | Bistable<br>Never in Alternating PD | $A_9 = \begin{pmatrix} 0.4 & 0.1 \\ 0.75 & 0.3 \end{pmatrix}$ |
| Genotype $G_3$ is not ESGD<br>$a_{22} < \frac{1}{5}(a_{11} + 3a_{12} + a_{21})$ | Never in Alternating PD | Coexistence<br>$A_{11} = \begin{pmatrix} 0.4 & 0.1 \\ 0.8 & 0.2 \end{pmatrix}$ |

Although coexistence does not guarantee the presence of an interior equilibrium, every numerical example in this article that satisfies the conditions for coexistence exhibits an (interior) ESGD, thereby demonstrating that Definition 1 is not empty. To illustrate this, let us consider the example of coexistence from Table 6. The payoff matrix is 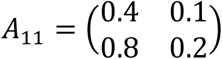. The corresponding interior Nash equilibrium, which is the solution to (6), is 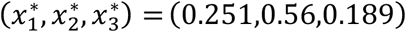. This equilibrium is evolutionarily stable, as shown in Figure 1.A, where the function *F*(*x*^*^, *x*) is plotted around *x*^*^, clearly indicating that *F*(*x*^*^, *x*) has a minimum in variable *x* at *x* = *x*^*^. Figure 1.B depicts the trajectories of the dynamics (1). Apparently, that the interior equilibrium point *x*^*^ is asymptotically stable, while the two trivial equilibrium points, (1,0,0) and (0,0,1), are unstable.

**Figure 1.**
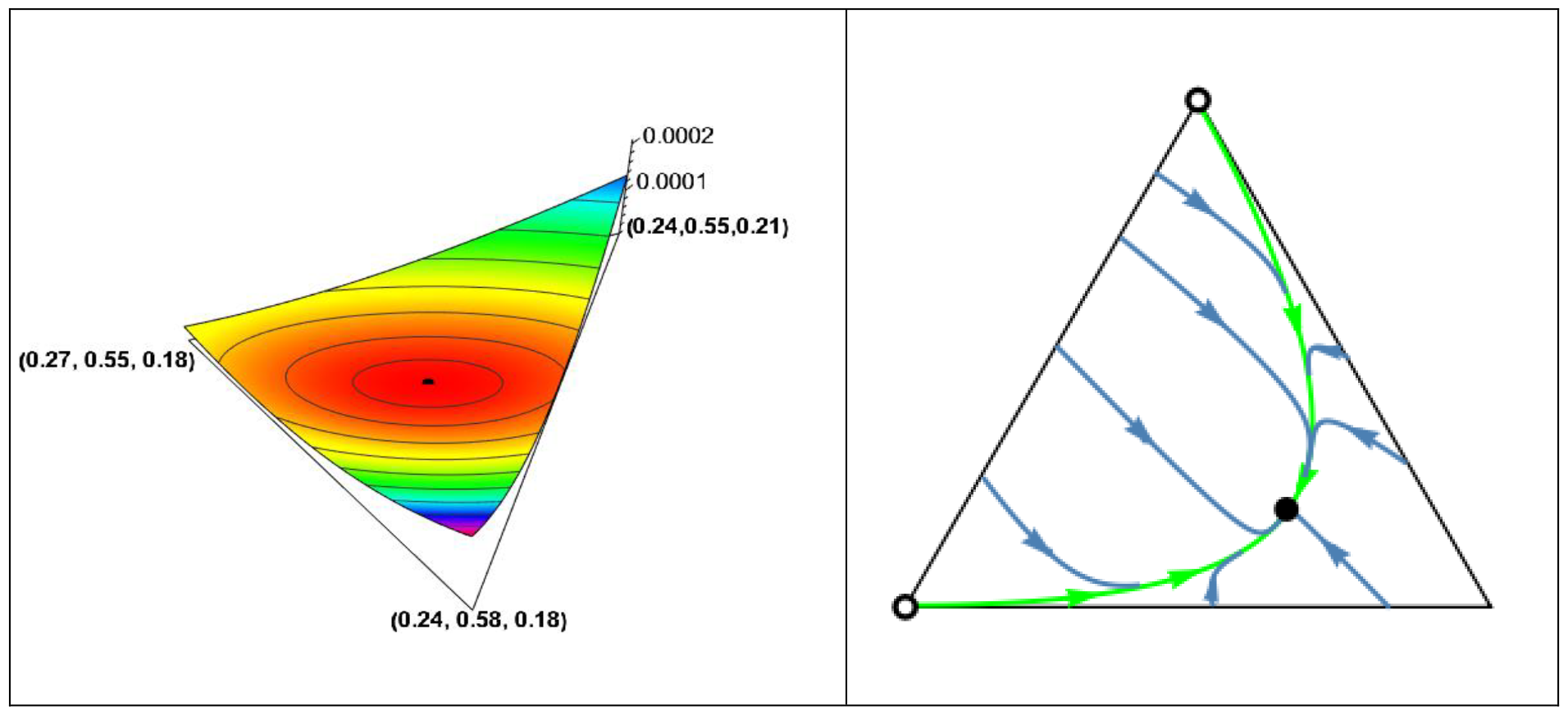
*Panel A*. Function *F*(*x*^*^, *x*) around *x*^*^ = (0.251,0.56,0.189). The black dot is the graph point (*x*^*^, *F*(*x*^*^, *x*^*^)) corresponding to *x*^*^. Clearly, that *F*(*x*^*^, *x*) in variable *x* has a minimum there. *Panel B*. Trajectories of dynamics (1). The system is governed by phenotypic payoff matrix 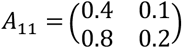 from Table 6. The globally asymptotically stable interior rest point is *x*^*^ = (0.251,0.56,0.189); 0.8 0.2 the homozygote vertices (1,0,0) and (0,0,1) are unstable rest points. Black dots and circles denote asymptotically stable and unstable rest point, respectively. Observe, the simplex *S*_3_ is only positively invariant under the genotype dynamics (1), in general. For instance, the homozygote vertices (1,0,0) and (0,0,1) are trivial rest points of the genotype dynamics (1), but the heterozygote vertex (0,1,0), is not a rest point. We note that the heterozygote vertex is a rest point of the genotype dynamics (1) if and only if both homozygotes are lethal.

## The cooperator phenotype is dominant

If the cooperator phenotype is dominant then *s*_1_ = *s*_2_ = (1,0) and *s*_3_ = (0,1). For the relating table, see Table 7.

Observe that Tables 4 and 7 illustrate different selection scenarios. In *G*_2_*×G*_2_ families, for instance, the survival rates vary. In particular, for each *k*, the value of *nk*(22) differs. According to the genotype-phenotype mapping, if the cooperator phenotype is dominant, there will be a higher number of cooperators in families with the same genotype distribution.

**Table 7.** Genotype survival table based on Table 1 if the cooperator strategy is dominant. The phenotype of genotype *G*_1_ and *G*_2_ is the cooperator strategy represented by *s*_1_ = (1,0), the phenotype of genotype *G*_3_ is the defector strategy represented by *s*_3_ = (0,1). *s*_1_ is dominant while *s*_3_ is recessive. *p*_*k*(*ij*)_ is the probability that the genotype of a given offspring is *G*_*k*_ provided the parental genotypes are *G*_*i*_ and *G*_*j*_ , respectively. *n*_*k*(*ij*)_ is the number of offspring with genotype *G*_*k*_ provided the parental genotypes are *G*_*i*_ and *G*_*j*_ , respectively. “*A*” is a payoff matrix describing the interactions between siblings.

| Genotypes of parents | Average number of couples assuming panmixia | Average number of survived family members, for genotypes $G_1, G_2, G_3$ | | |
| --- | --- | --- | --- | --- |
| | | $G_1 \equiv ([a],[a])$<br>dominant $s_1 = (1,0)$ | $G_2 \equiv ([a],[A])$<br>dominant $s_2 = s_1 = (1,0)$ | $G_3 \equiv ([A],[A])$,<br>recessive $s_3 = (0,1)$ |
| $G_1 \times G_1$ | $\frac{N}{2} x_1^2$ | $n_{1(11)} = na_{11}$ | 0 | 0 |
| $G_1 \times G_2$ | $Nx_1x_2$ | $n_{1(12)} = \frac{n}{2}a_{11}$ | $n_{2(12)} = \frac{n}{2}a_{11}$ | 0 |
| $G_1 \times G_3$ | $Nx_1x_3$ | 0 | $n_{2(13)} = na_{11}$ | 0 |
| $G_2 \times G_2$ | $\frac{N}{2} x_2^2$ | $n_{1(22)} = \frac{n}{16}(3a_{11} + a_{12})$ | $n_{2(22)} = \frac{n}{8}(3a_{11} + a_{12})$ | $n_{3(22)} = \frac{n}{16}(3a_{21} + a_{22})$ |
| $G_2 \times G_3$ | $Nx_2x_3$ | 0 | $n_{2(23)} = \frac{n}{4}(a_{11} + a_{12})$ | $n_{3(23)} = \frac{n}{4}(a_{21} + a_{22})$ |
| $G_3 \times G_3$ | $\frac{N}{2} x_3^2$ | 0 | 0 | $n_{3(33)} = na_{22}$ |

If allele [a] of the cooperation is dominant, then the condition of the stability of the homozygous states reads as follows (see SI C).

State (1,0,0), that is, the state in which every individual is *G*_1_ = ([*a*], [*a*]) homozygote, is a local ESGD if

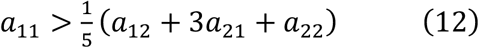

State (0,0,1), that is the state in which every individual is *G*_3_ = ([*A*], [*A*]) homozygote, is a local ESGD if

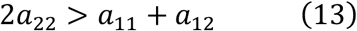

Similarly to the recessive case, if we reverse the inequality in (12), it ensures that state (1,0,0) is not an ESGD. Likewise, reversing the inequality in (13) implies that state (0,0,1) is not an ESGD. It should be noted that conditions (12–13) apply to arbitrary payoff matrices when phenotype *s*_1_ is dominant. When the prisoner’s dilemma describes the interactions, based on the stability of (1,0,0) and (0,0,1), we can again identify four subcases in the collaborating case and two subcases in the alternating case. These subcases are illustrated with particular examples in Tables 8 and 9.

**Table 8.** Examples where cooperation is dominant in collaborating PD game. Black dots correspond to an asymptotically stable rest points while empty dots represent unstable equilibrium points. The bottom-left vertex of the triangle corresponds to the (1,0,0) state, in which all individuals are ([a],[a]) cooperator homozygotes. The bottom-right vertex corresponds to the (0,1,0) state, where all individuals are ([a],[A]) heterozygotes. The top vertex corresponds to the (0,0,1) state, where all individuals are ([A],[A]) defector homozygotes. Black dots and circles denote asymptotically stable and unstable rest point, respectively.

| Collaborate<br>$2a_{11} > a_{12} + a_{21}$ | Genotype $G_1$ is ESGD<br>$a_{11} > \frac{1}{5}(a_{12} + 3a_{21} + a_{22})$ | Genotype $G_1$ is not ESGD<br>$a_{11} < \frac{1}{5}(a_{12} + 3a_{21} + a_{22})$ |
| --- | --- | --- |
| Genotype $G_3$ is ESGD<br>$2a_{22} > a_{11} + a_{12}$ | Bistable<br>$A_1 = \begin{pmatrix} 0.6 & 0.1 \\ 0.7 & 0.4 \end{pmatrix}$ | $A_7 = \begin{pmatrix} 0.3 & 0.1 \\ 0.49 & 0.21 \end{pmatrix}$ |
| Genotype $G_3$ is not ESGD<br>$2a_{22} < a_{11} + a_{12}$ | $A_3 = \begin{pmatrix} 0.7 & 0.1 \\ 0.8 & 0.2 \end{pmatrix}$ | Coexistence<br>$A_4 = \begin{pmatrix} 0.5 & 0.1 \\ 0.8 & 0.15 \end{pmatrix}$ |

**Table 9.**
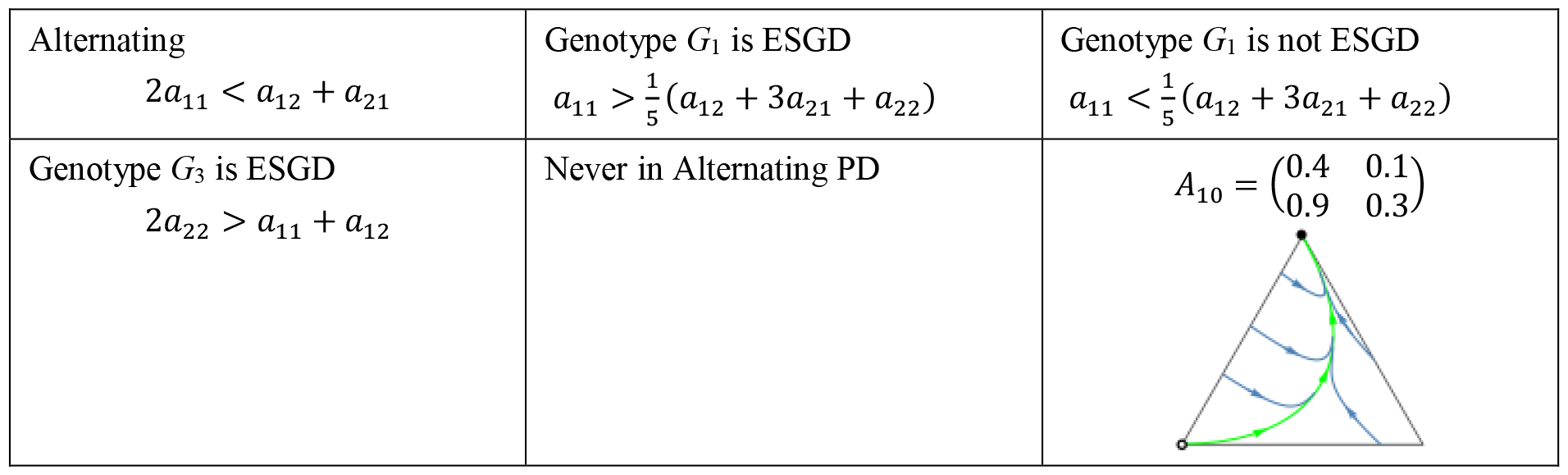
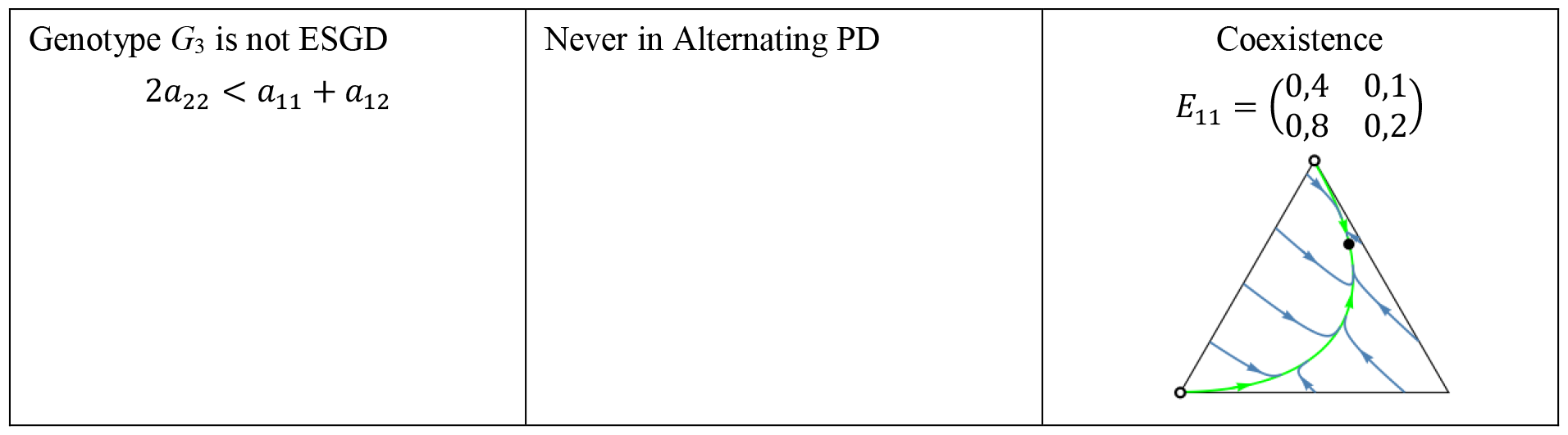
Examples where cooperation is recessive in alternating PD game. See the caption at Table 8.

| Alternating<br>$2a_{11} < a_{12} + a_{21}$ | Genotype $G_1$ is ESGD<br>$a_{11} > \frac{1}{5}(a_{12} + 3a_{21} + a_{22})$ | Genotype $G_1$ is not ESGD<br>$a_{11} < \frac{1}{5}(a_{12} + 3a_{21} + a_{22})$ |
| --- | --- | --- |
| Genotype $G_3$ is ESGD<br>$2a_{22} > a_{11} + a_{12}$ | Never in Alternating PD | $A_{10} = \begin{pmatrix} 0.4 & 0.1 \\ 0.9 & 0.3 \end{pmatrix}$ |
| Genotype $G_3$ is not ESGD<br>$2a_{22} < a_{11} + a_{12}$ | Never in Alternating PD | Coexistence<br>$E_{11} = \begin{pmatrix} 0,4 & 0,1 \\ 0,8 & 0,2 \end{pmatrix}$ |

## Intermediate inheritance

Finally, we turn to the analysis of intermediate inheritance. In this case, the phenotype of the *G*_1_ = ([*a*], [*a*]) homozygotes is the pure strategy (1,0), the phenotype of the *G*_3_ = ([*A*], [*A*]) homozygotes is the pure strategy (0,1), while the phenotype of the *G*_2_ = ([*a*], [*A*]) heterozygotes is the mixed strategy (1*⁄*2 , 1*⁄*2) (see Table 10).

**Table 10.** Genotype survival table based on Table 1 in case of intermediate inheritance. The phenotype of genotype *G*_1_ is the cooperator strategy represented by *s*_1_ = (1,0), the phenotype of genotype *G*_3_ is the defector strategy represented by *s*_3_ = (0,1) while the phenotype of the heterozygote *G*_2_ is (1*⁄*2,1*⁄*2) denoted by *s*_2_. *p*_*k*(*ij*)_ is the probability that the genotype of a given offspring is *G*_*k*_ provided the parental genotypes are *G*_*i*_ and *G*_*j*_ , respectively. *n*_*k*(*ij*)_ is the number of offspring with genotype *G*_*k*_ provided the parental genotypes are *G*_*i*_ and *G*_*j*_ , respectively. “*A*” is a payoff matrix describing the interactions between siblings.

| Genotypes of parents | Average number of couples<br>$\frac{N}{2} h_{ij}(x)$ | Average number of survived family members, for genotypes $G_1, G_2, G_3$ | | |
| --- | --- | --- | --- | --- |
| | | $G_1 \equiv ([a], [a])$<br>$s_1 = (1,0)$ | $G_2 \equiv ([a], [A])$<br>$s_2 = (\frac{1}{2}, \frac{1}{2})$ | $G_3 \equiv ([A], [A])$<br>$s_3 = (0,1)$ |
| $G_1 \times G_1$ | $\frac{N}{2} x_1^2$ | $n_{1(11)} = na_{11}$ | 0 | 0 |
| $G_1 \times G_2$ | $Nx_1x_2$ | $n_{1(12)} = \frac{n}{8}(3a_{11} + a_{12})$ | $n_{2(12)} = \frac{n}{16}(3a_{11} + a_{12} + 3a_{21} + a_{22})$ | 0 |
| $G_1 \times G_3$ | $Nx_1x_3$ | 0 | $n_{2(13)} = \frac{n}{4}(a_{11} + a_{12} + a_{21} + a_{22})$ | 0 |
| $G_2 \times G_2$ | $\frac{N}{2} x_2^2$ | $n_{1(22)} = \frac{n}{8}(a_{11} + a_{12})$ | $n_{2(22)} = \frac{n}{8}(a_{11} + a_{12} + a_{21} + a_{22})$ | $n_{3(22)} = \frac{n}{8}(a_{21} + a_{22})$ |
| $G_2 \times G_3$ | $Nx_2x_3$ | 0 | $n_{2(23)} = \frac{n}{16}(a_{11} + 3a_{12} + a_{21} + 3a_{22})$ | $n_{3(23)} = \frac{n}{8}(a_{21} + 3a_{22})$ |
| $G_3 \times G_3$ | $\frac{N}{2} x_3^2$ | 0 | 0 | $n_{3(33)} = na_{22}$ |

Similarly to the recessive and the dominant cases, we give a mathematical conditions for the stability of the homozygote states (1,0,0) and (0,0,1). In the main text, we only mention the first order condition. The detailed mathematical answer is in SI C.

State (1,0,0), that is, the state in which every individual is a *G*_1_ = ([*a*], [*a*]) homozygote, is a local ESGD if

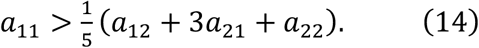

State (0,0,1), that is, the state in which every individual is a *G*_3_ = ([*A*], [*A*]) homozygote, is a local ESGD if

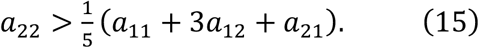

Furthermore, if we reverse the inequality in (14), the state (1,0,0) is no longer an ESGD, and if we reverse the inequality in (15), the state (0,0,1) is not an ESGD either. It’s important to note that these conclusions hold true for arbitrary matrix games under intermediate inheritance.

When we analyze the stability of the homozygous states (1,0,0) and (0,0,1) in the genotype dynamics using inequalities (14–15) in the context of prisoner’s dilemma, we once again have four possible subcases in the collaborating case and two possible subcases in the alternating case. These scenarios are illustrated with concrete numerical examples in Tables 11 and 12.

**Table 11.**
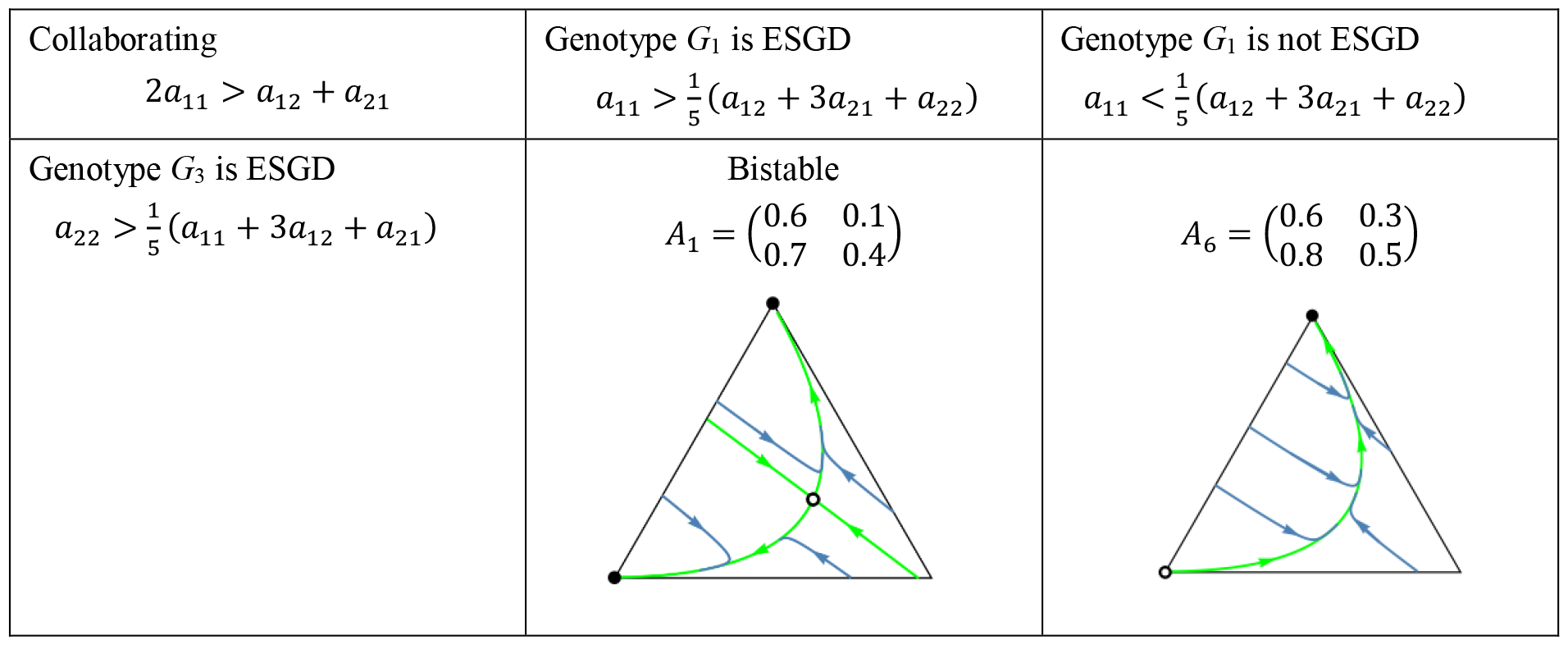
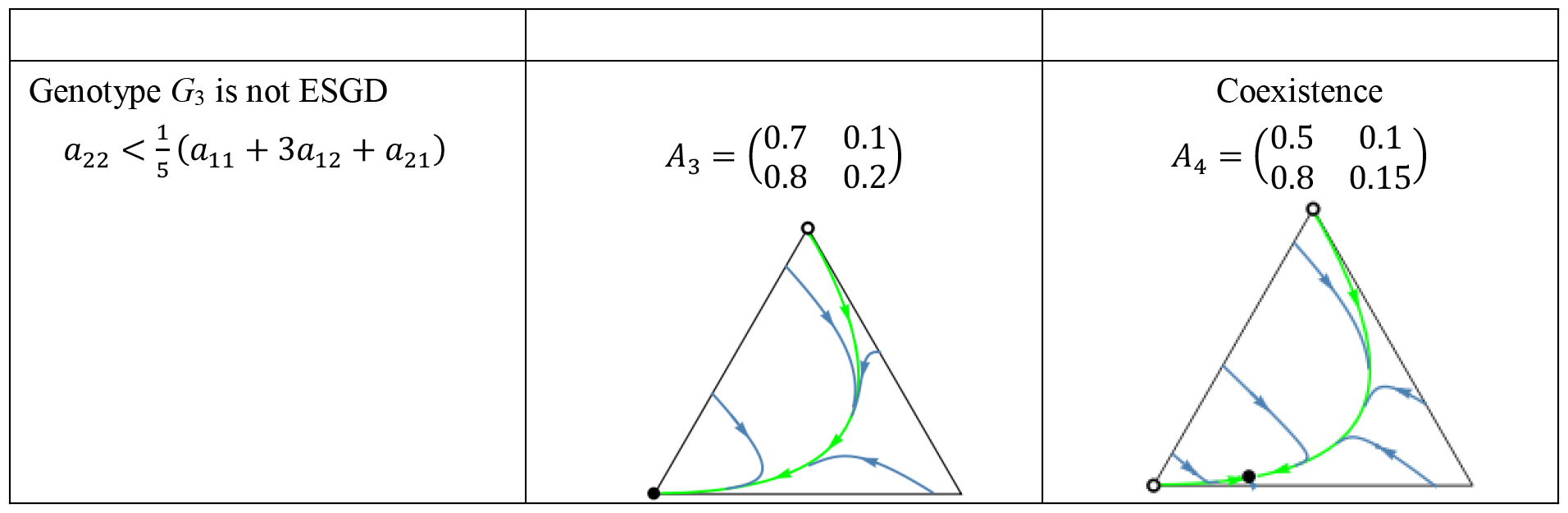
Examples in case of intermediate inheritance in collaborating PD game. Black dots correspond to an asymptotically stable rest points while empty dots represent unstable equilibrium points. The bottom-left vertex of the triangle corresponds to the (1,0,0) state, in which all individuals are ([a],[a]) cooperator homozygotes. The bottom-right vertex corresponds to the (0,1,0) state, where all individuals are ([a],[A]) heterozygotes. The top vertex corresponds to the (0,0,1) state, where all individuals are ([A],[A]) defector homozygotes. Black dots and circles denote asymptotically stable and unstable rest point, respectively.

| Collaborating<br>$2a_{11} > a_{12} + a_{21}$ | Genotype $G_1$ is ESGD<br>$a_{11} > \frac{1}{5}(a_{12} + 3a_{21} + a_{22})$ | Genotype $G_1$ is not ESGD<br>$a_{11} < \frac{1}{5}(a_{12} + 3a_{21} + a_{22})$ |
| --- | --- | --- |
| Genotype $G_3$ is ESGD<br>$a_{22} > \frac{1}{5}(a_{11} + 3a_{12} + a_{21})$ | Bistable<br>$A_1 = \begin{pmatrix} 0.6 & 0.1 \\ 0.7 & 0.4 \end{pmatrix}$ | $A_6 = \begin{pmatrix} 0.6 & 0.3 \\ 0.8 & 0.5 \end{pmatrix}$ |
| Genotype $G_3$ is not ESGD<br>$a_{22} < \frac{1}{5}(a_{11} + 3a_{12} + a_{21})$ | $A_3 = \begin{pmatrix} 0.7 & 0.1 \\ 0.8 & 0.2 \end{pmatrix}$ | Coexistence<br>$A_4 = \begin{pmatrix} 0.5 & 0.1 \\ 0.8 & 0.15 \end{pmatrix}$ |

**Table 12.**
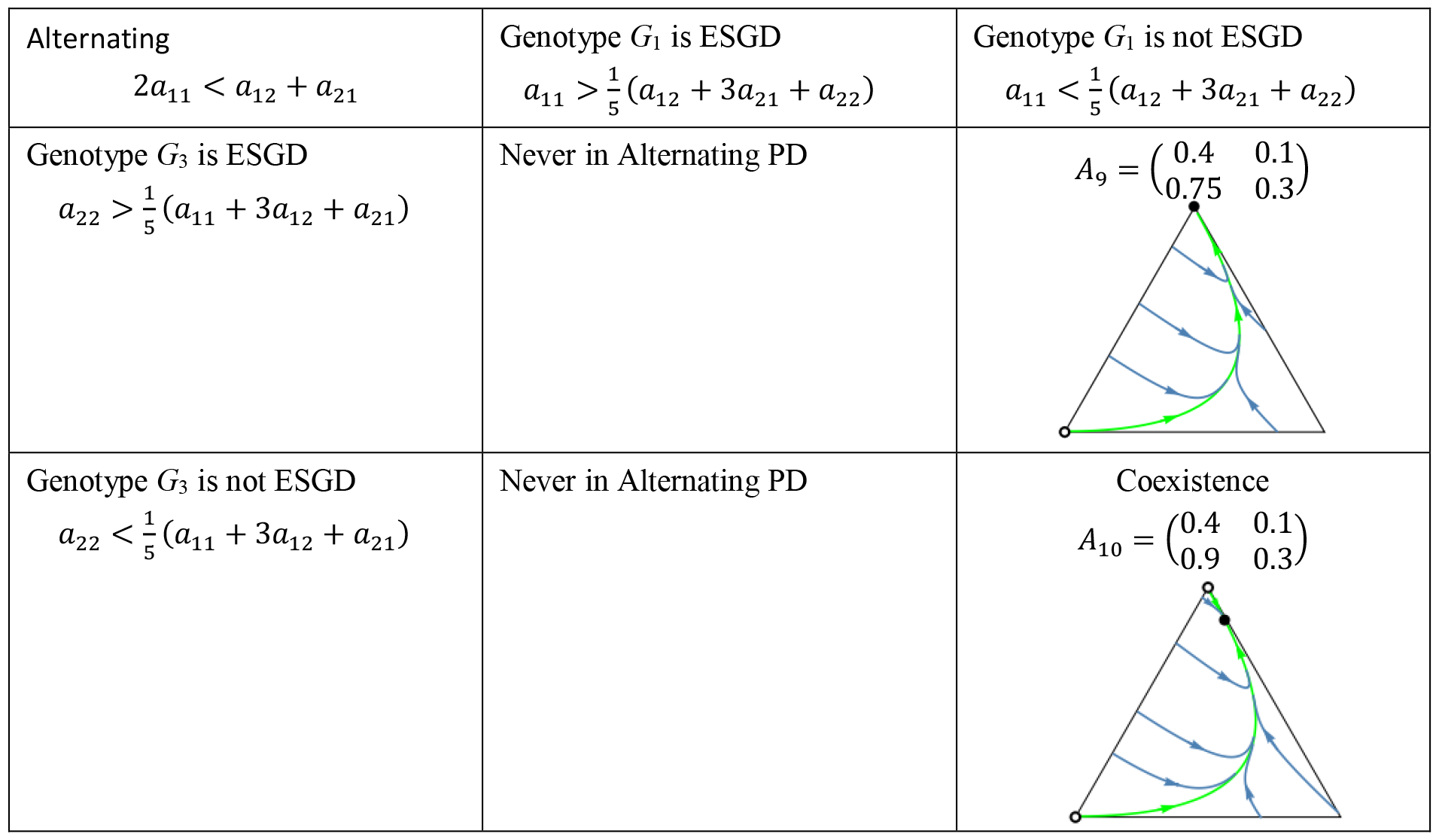
Examples in case of intermediate inheritance in alternating PD game. See the caption at Table 11.

|  |  |  |
| --- | --- | --- |
| Alternating<br>$2a_{11} < a_{12} + a_{21}$ | Genotype $G_1$ is ESGD<br>$a_{11} > \frac{1}{5}(a_{12} + 3a_{21} + a_{22})$ | Genotype $G_1$ is not ESGD<br>$a_{11} < \frac{1}{5}(a_{12} + 3a_{21} + a_{22})$ |
| Genotype $G_3$ is ESGD<br>$a_{22} > \frac{1}{5}(a_{11} + 3a_{12} + a_{21})$ | Never in Alternating PD | $A_9 = \begin{pmatrix} 0.4 & 0.1 \\ 0.75 & 0.3 \end{pmatrix}$ |
| Genotype $G_3$ is not ESGD<br>$a_{22} < \frac{1}{5}(a_{11} + 3a_{12} + a_{21})$ | Never in Alternating PD | Coexistence<br>$A_{10} = \begin{pmatrix} 0.4 & 0.1 \\ 0.9 & 0.3 \end{pmatrix}$ |

## Donation game in diploid Mendelian population and classical Hamilton’s rule

A special case of prisoner’s dilemma is the so-called “donation game” [20], with the payoff matrix

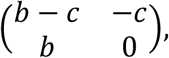

where *b* and *c* are the benefit and the cost of an interaction, respectively, with *b* > *c*. Donation game can be considered as a connection between the prisoner’s dilemma and Hamilton’s rule [25-27]. Since in monogamous diploid family the relatedness between full siblings is *r* = 1/2, the classical Hamilton’s rule predicts cooperation between siblings when *b* > 2*c*.

The donation game is not a survival game (the entries of the payoff matrix are not survival probabilities), although starvation can be one of the main causes of death. In case of famine, when survival requires reaching a minimum level of food intake during the critical period, siblings can help each other through food sharing. Therefore, an additive model (where the outcomes of individual games are summed) is more suitable for modeling food sharing than a multiplicative model (where the outcomes of individual games are multiplied).

We note that only non-negative payoff matrices make sense in the model. However, it is not a limitation because if we add the same number to every element of the matrix, the relative fitness and therefore the game-theoretical predictions remain unchanged. Therefore, instead of using the above payoff matrix, we can use the following matrix:

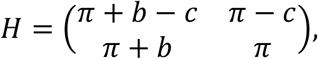

where *π* > *c* can be interpreted as an interaction-independent basic fitness.

Now the question arises: what condition ensures the local stability of the cooperator homozygotes *G*_1_ in the game between diploid siblings? Since *b* > *c*, the payoff matrix *H* satisfies *a*_21_ > *a*_11_ > *a*_22_ > *a*_12_ and 2*a*_11_ > *a*_21_ *+ a*_12_, indicating that the donation game represents a specific version of collaborating prisoner’s dilemma games. It is easy to see, that inequalities (8), (12) and (14) can be read as the classical Hamilton’s rule, while inequalities (9), (13), and (15) are equivalent to *b <* 2*c*. Therefore, within the considered inheritance systems where the donation game determines the survival rate of siblings, there is neither bistability nor coexistence. There are only two possibilities here: if *b* > 2*c* then cooperator homozygotes G_1_, whereas if *b <* 2*c* then defector homozygotes G_3_ become fixed through selection.

## Haldane monotonicity

Price [28] was interested in how the relative frequency of an allele changes between the parental and the offspring population. Hamilton’s rule [27, 29] and Haldane’s arithmetic [4] describe conditions under which the frequency of the altruistic allele strictly increases in well-defined selection scenario, ensuring the evolutionary stability of the population of altruistic homozygotes. (Haldane’s arithmetic claims [4]: “*The relative frequency of the altruistic gene increases, if by ‘self-sacrifice’, it has ‘rescued’ on average more than one copy of itself in its lineage*.” (We note that Haldane’s arithmetic is equivalent with evolutionary stability (in ESGD sense) of recessive homozygotes in the altruistic one-person game. Moreover, this equivalence holds even when the interaction between siblings is described by a non-linear game [4].) These rules are effective when fitness can be expressed using explicit cost and benefit parameters. In the case of a prisoner’s dilemma game, however, at least three parameters are typically required, making the application of both Hamilton’s rule and Haldane’s arithmetic less straightforward.

Instead, we are seeking a different approach to answer when the frequency of an allele increases along every trajectory of the genotype dynamics, or mathematically speaking, when the frequency of the given allele acts as a Lyapunov function. For simplicity, we consider only the case where three genotypes are possible. Consider the frequency of the allele in homozygote G_1_ at a genotype distribution *x* ∈ *S*_3_. It is

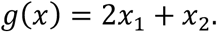

The derivative of function *g* with respect to the genotype dynamics (1) is

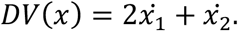

Since for the genotype dynamics we have 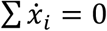, it follows that 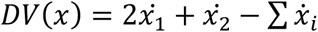, so

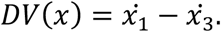

Thus, the frequency of the allele in homozygote G_1_ is strictly increases if

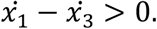

The biological interpretation of this inequality is as follows: Let us refer to genotype dynamics as Haldane monotonic if the relative advantage of the *G*_1_ homozygote over the entire population is greater than that of the *G*_3_ homozygote. The key distinction between Hamilton’s rule, Haldane’s arithmetic, and Haldane monotonicity lies in their respective focuses. Hamilton’s rule emphasizes the interaction between two siblings, Haldane’s arithmetic examines how many parental genes are saved through the “self-sacrifice” of a cooperating sibling, while Haldane monotonicity centers on the relative advantage of a particular homozygote and also takes into account the production rates within the entire population.

### Remark 4.

Mathematically, Haldane monotonicity provides only a sufficient condition for global stability of a homozygous state, as global stability can still occur even if Haldane monotonicity is not satisfied. For example, it is possible that one trajectory initially moves away from the homozygous vertex but then reverses (cf. Figure 3.).

Firstly, we illustrate through a numerical example the existence of a matrix game where the genotype dynamics is Haldane monotone, see Figure 2.

**Figure 2.**
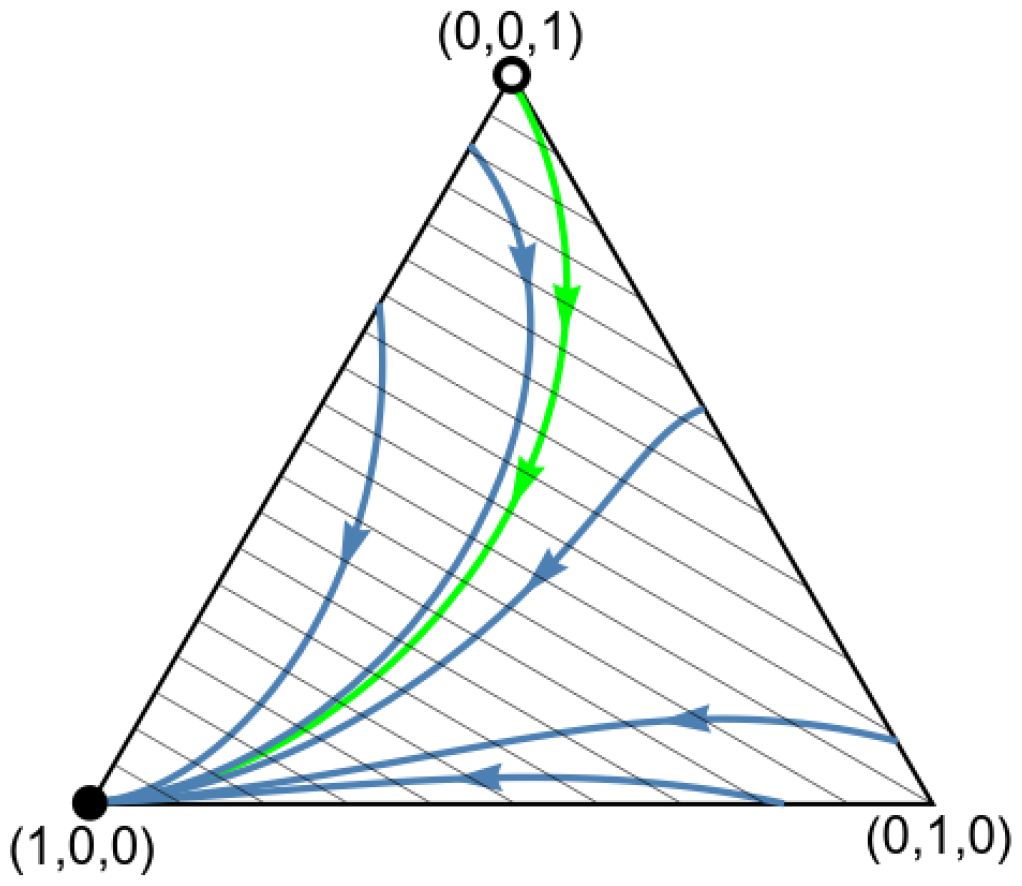
Here we consider the payoff matrix 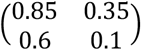 and let gene [a] be recessive under [A]. Notice that the orbits, while moving toward the vertex (1,0,0) corresponding to the recessive homozygotes (that is the genotype of every individual is ([a],[a])), intersect the level lines (which are the parallel lines in the figure) of the function 2*x*_1_ *+ x*_2_, which describes the frequency of the recessive allele, exactly once. Vertex (0,1,0) is the state in which is the genotype of every individual is ([a],[A]) while vertex (0,0,1) is the state in which is the genotype of every individual is ([A],[A]). The black dot corresponds to an asymptotically stable rest point while the empty dot represents an unstable equilibrium point.

Secondly, the question arises: Is the genotype dynamic Haldane monotone when the donation game takes place between siblings? The answer is negative, see Figure 3. The intuitive reason for that is the following. When the whole population contains only heterozygotes, according to Mendelian inheritance, the diploid embryos follow Hardy-Weinberg proportions, i.e. 1/4, 1/2 and 1/4 are the respective relative frequencies of the recessive homozygote, heterozygote and dominant homozygote. If the phenotypic selection cannot change radically these Hardy-Weinberg proportions, then the frequency of the cooperative allele cannot increase. Here the main point is that the genetic system and the phenotypic selection together determine the frequency change of genotypes. We note the heterozygote vertex is a rest point of the genotype dynamics only if both homozygotes are lethal in all families.

**Figure 3.**
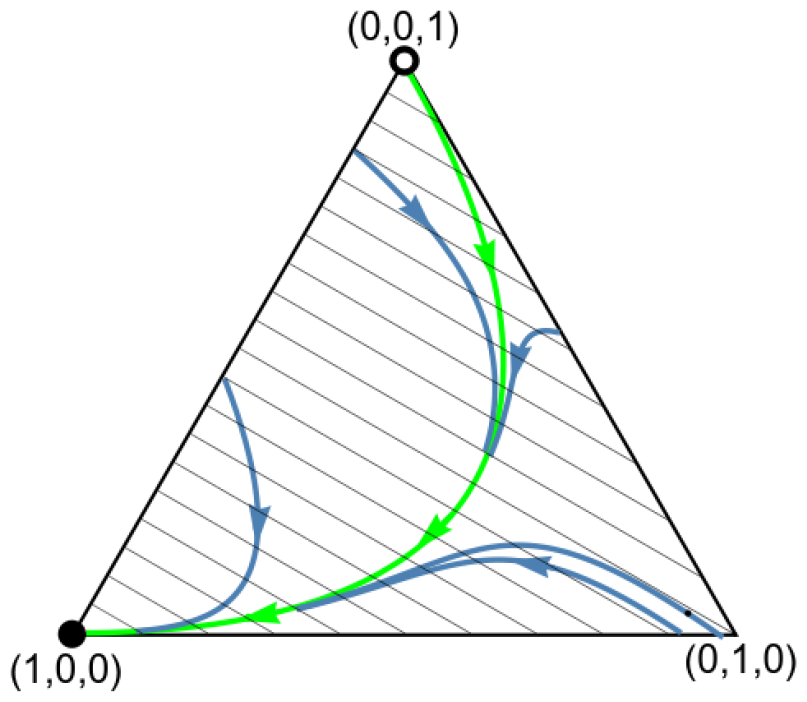
Consider a donation game with *π* = 0.2, *b* = 0.6 and *c* = 0.1, thus the payoff matrix reads 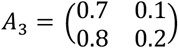 and let allele [a] be recessive under allele [A]. The parallel lines are the level lines of the function 2*x*_1_ *+ x*_2_, which describes the frequency of recessive alleles. Observe that the function first decreases, then increases along the orbits starting near the heterozygote vertex (0,1,0) (which is not an equilibrium). This shows that this dynamics globally is not Haldane monotone. However, if we discount an appropriate neighborhood of the pure heterozygote vertex, the system is Haldane monotone. Particularly, at the neighborhood of the pure recessive ([a],[a]) homozygote vertex (1,0,0), the system is locally Haldane monotone. Vertex (0,0,1) is the pure dominant ([A],[A]) homozygote vertex in the sense that the genotype of every individual is ([A],[A]). The black dot corresponds to an asymptotically stable rest point while the empty dot represents an unstable equilibrium point.

## Discussion

### In diploid population, the payoff matrix and the genotype-phenotype mapping together determine the endpoint of the evolution

Firstly, we examine the conditions under which the homozygous states are evolutionarily stable, depending on the inheritance system and the (arbitrary) survival payoff matrix (i.e., where each matrix element is between 0 and 1). Applying the results in SI.B in [3] for matrix games, we obtain the following Table 13.

**Table 13.** Summary of the condition for arbitrary payoff matrices (*a*_*ij*_ )_2*×*2_. [a] and [A] are different alleles of the same gene. (1,0,0) is the state when the genotype of every individual is *G*_1_=([a],[a]) with phenotype (1,0). (0,0,1) is the state when the genotype of every individual is *G*_3_=([A],[A]) with phenotype (0,1). [a] and [A] are different alleles of the same gene. (*a*_*ij*_ )_2*×*2_ is a 2×2 payoff matrix describing the interactions between siblings.

| | Genotype $G_1 = ([a],[a])$ is ESGD | Genotype $G_3 = ([A],[A])$ is ESGD |
| --- | --- | --- |
| [a] is recessive | $2a_{11} > a_{21} + a_{22}$ | $a_{22} > \frac{1}{5}(a_{11} + 3a_{12} + a_{21})$ |
| [a] is dominant | $a_{11} > \frac{1}{5}(a_{12} + 3a_{21} + a_{22})$ | $2a_{22} > a_{11} + a_{12}$ |
| Intermediate inheritance | $a_{11} > \frac{1}{5}(a_{12} + 3a_{21} + a_{22})$ | $a_{22} > \frac{1}{5}(a_{11} + 3a_{12} + a_{21})$ |

Note the following observations:

Note the following observations:

- The condition for dominant *G*_3_ is the same as the condition for intermediate *G*_3_, hence in the case of recessive and intermediate inheritance, the state associated with *G*_3_ is simultaneously ESS or non-ESS or degenerate.
- The condition for dominant *G*_1_ is the same as the condition for intermediate *G*_1_, therefore, in the case of dominant and intermediate inheritance, the state associated with *G*_1_ is simultaneously ESS or non-ESS or degenerate.
- Consequently, the recessive and dominant cases (with respect to the allele in *G*_1_) unambiguously determine what happens in the intermediate case. In particular, in the intermediate case, the stability of the state associated with *G*_1_ is the same as in the dominant case, while the stability of the state associated with *G*_3_ is the same as in the recessive case.

We note these coincidences are surprising, since the selection regimes given by Tables 4, 7 and 10 are quite different.

Secondly, based on the previous observations, we can classify prisoner’s dilemma matrices according to the stability of the two homozygous states under different inheritance systems. We find that in the case of collaborating matrices, there are 8 possible scenarios, while in the case of alternating matrices, there are only 3. These are summarized in Table 14 and 15 accompanied by numerical examples illustrating each case.

**Table 14.**
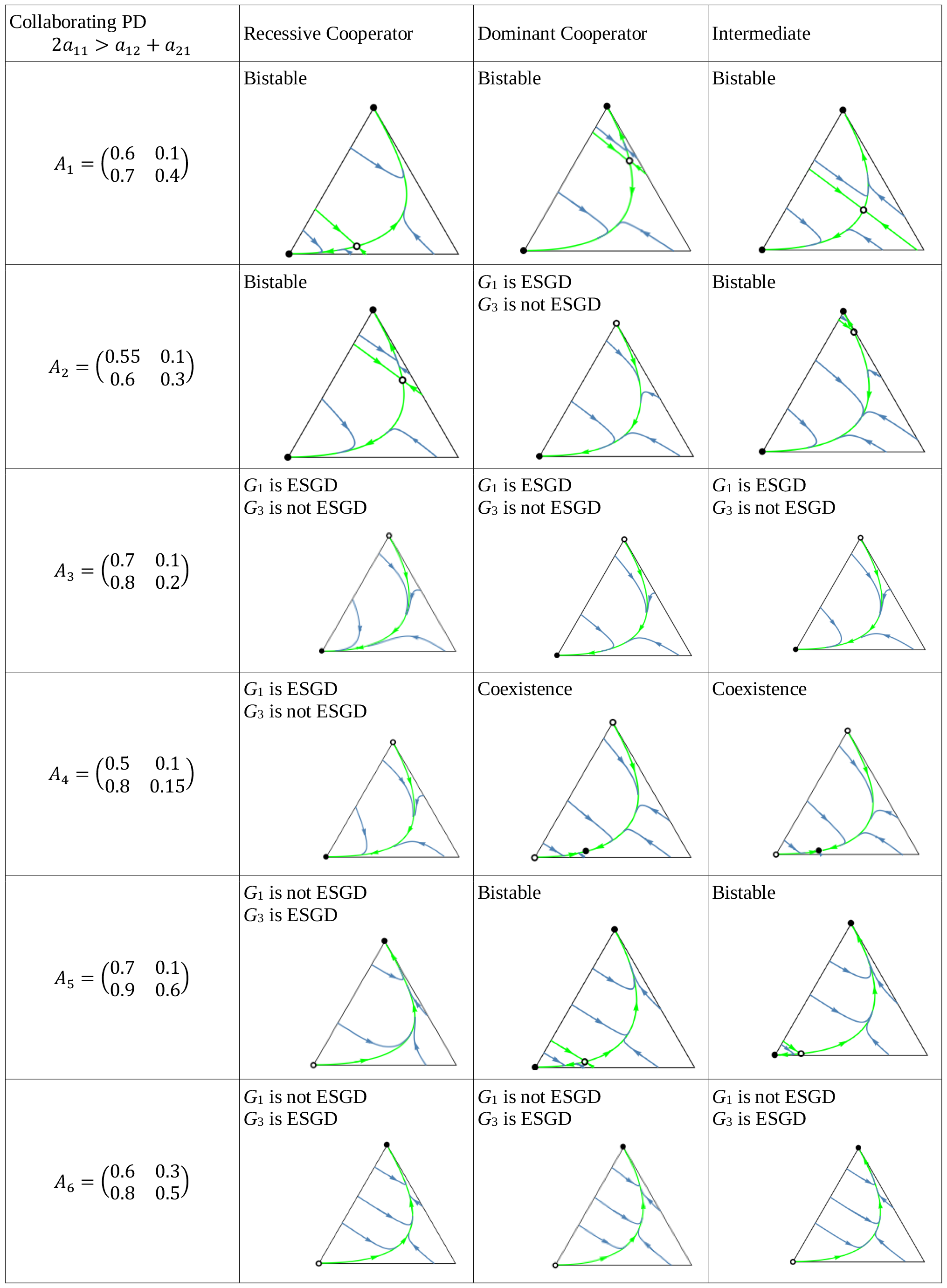
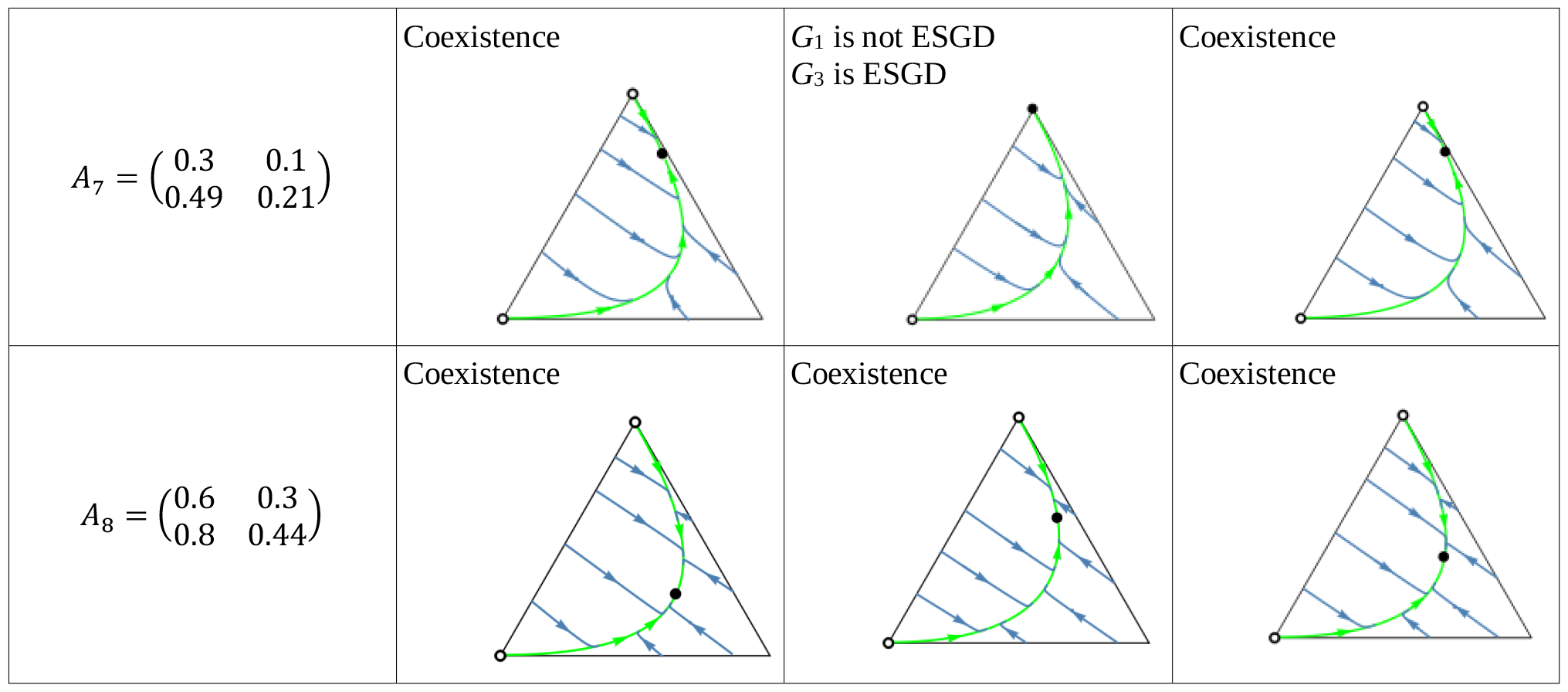
Summary of collaborating PD. In the right column, the payoff matrices are given. In the other columns, the phase portraits of the corresponding genotype dynamics (1). Black dots correspond to a asymptotically stable rest points while empty dots represent unstable equilibrium points. The bottom-left vertex of the triangle corresponds to the (1,0,0) state, in which all individuals are ([a],[a]) cooperator homozygotes. The bottom-right vertex corresponds to the (0,1,0) state, where all individuals are ([a],[A]) heterozygotes. The top vertex corresponds to the (0,0,1) state, where all individuals are ([A],[A]) defector homozygotes.

| Collaborating PD<br>$2a_{11} > a_{12} + a_{21}$ | Recessive Cooperator | Dominant Cooperator | Intermediate |
| --- | --- | --- | --- |
| $A_1 = \begin{pmatrix} 0.6 & 0.1 \\ 0.7 & 0.4 \end{pmatrix}$ | Bistable | Bistable | Bistable |
| $A_2 = \begin{pmatrix} 0.55 & 0.1 \\ 0.6 & 0.3 \end{pmatrix}$ | Bistable | $G_1$ is ESGD<br>$G_3$ is not ESGD | Bistable |
| $A_3 = \begin{pmatrix} 0.7 & 0.1 \\ 0.8 & 0.2 \end{pmatrix}$ | $G_1$ is ESGD<br>$G_3$ is not ESGD | $G_1$ is ESGD<br>$G_3$ is not ESGD | $G_1$ is ESGD<br>$G_3$ is not ESGD |
| $A_4 = \begin{pmatrix} 0.5 & 0.1 \\ 0.8 & 0.15 \end{pmatrix}$ | $G_1$ is ESGD<br>$G_3$ is not ESGD | Coexistence | Coexistence |
| $A_5 = \begin{pmatrix} 0.7 & 0.1 \\ 0.9 & 0.6 \end{pmatrix}$ | $G_1$ is not ESGD<br>$G_3$ is ESGD | Bistable | Bistable |
| $A_6 = \begin{pmatrix} 0.6 & 0.3 \\ 0.8 & 0.5 \end{pmatrix}$ | $G_1$ is not ESGD<br>$G_3$ is ESGD | $G_1$ is not ESGD<br>$G_3$ is ESGD | $G_1$ is not ESGD<br>$G_3$ is ESGD |
| $A_7 = \begin{pmatrix} 0.3 & 0.1 \\ 0.49 & 0.21 \end{pmatrix}$ | Coexistence | $G_1$ is not ESGD<br>$G_3$ is ESGD | Coexistence |
| $A_8 = \begin{pmatrix} 0.6 & 0.3 \\ 0.8 & 0.44 \end{pmatrix}$ | Coexistence | Coexistence | Coexistence |

**Table 15.**
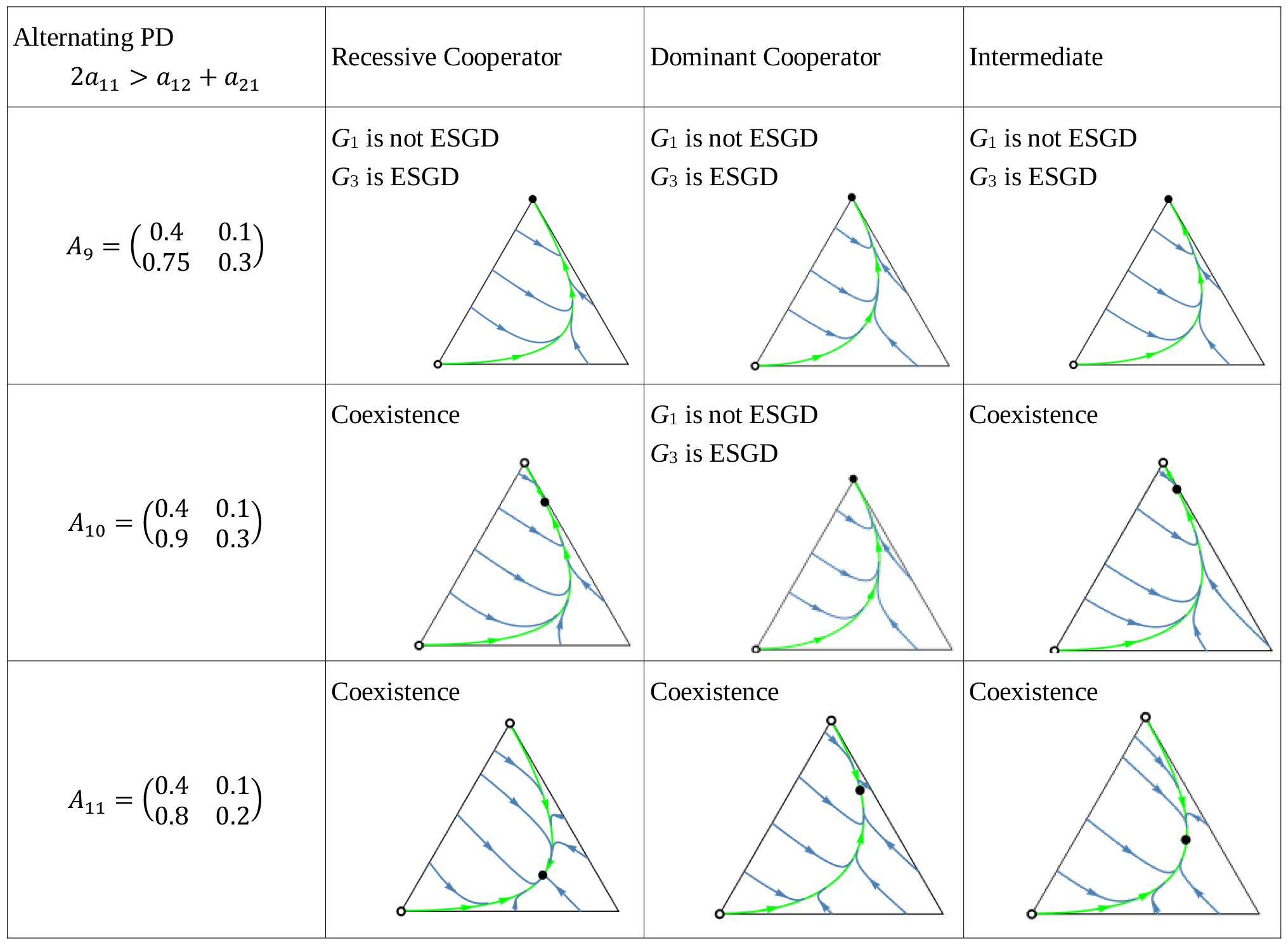
Summary of alternating PD See the caption at Table 14.

| Alternating PD<br>$2a_{11} > a_{12} + a_{21}$ | Recessive Cooperator | Dominant Cooperator | Intermediate |
| --- | --- | --- | --- |
| $A_9 = \begin{pmatrix} 0.4 & 0.1 \\ 0.75 & 0.3 \end{pmatrix}$ | $G_1$ is not ESGD<br>$G_3$ is ESGD | $G_1$ is not ESGD<br>$G_3$ is ESGD | $G_1$ is not ESGD<br>$G_3$ is ESGD |
| $A_{10} = \begin{pmatrix} 0.4 & 0.1 \\ 0.9 & 0.3 \end{pmatrix}$ | Coexistence | $G_1$ is not ESGD<br>$G_3$ is ESGD | Coexistence |
| $A_{11} = \begin{pmatrix} 0.4 & 0.1 \\ 0.8 & 0.2 \end{pmatrix}$ | Coexistence | Coexistence | Coexistence |

The numerical examples presented in Tables 14 and 15 call the attention for the followings:

- The cooperator and defector homozygote states can be fixed by natural selection, see e.g. *A*_3_ and *A*_6_, respectively.
- There are bistable cases (e.g. *A*_1_ ) and there is only one unstable interior rest point in such cases.
- The cooperator and defector genotypes can coexist (e.g. *A*_8_), and there is only one globally stable interior rest point in such cases. The genotype-phenotype map can change the coordinates of the stable rest point of the genotype dynamics.
- Although the phenotypic payoff matrix alone is not sufficient to predict the fixation of cooperation, it plays a crucial role in determining the outcome, since, in alternating prisoner’s dilemma, cooperator homozygotes cannot be fixed, but it is possible for cooperators to coexist with defectors see *A*_11_.
- Despite the 1/2 genetic relatedness between siblings and the fact that cooperation maximizes the overall survival rate of siblings in each collaborating prisoner’s dilemma, it is still possible for the defector homozygote to become fixed (e.g. *A*_6_).
- The phenotypic payoff matrix and the genotype-phenotype mapping together determine the end of the natural selection. When the phenotypic payoff matrix is fixed, the genotype dynamics can not only change the coordinates of the stable rest point, but also affect predictions regarding the presence of homozygotes (e.g. *A*_10_).
- However, note that there is no case where one inheritance system exhibits bistability while another exhibits coexistence; at most, only the stability of one homozygous vertex can change. This indicates that although the genotype-phenotype mapping can influence the outcome of natural selection, the payoff matrix still constrains evolution to some extent.

### Our results from the viewpoint of group selection

During the classification of prisoner’s dilemma games, the principle of “family welfare” has proven to be important, which means that siblings aim at maximizing the sum of their survival rates. We found that this principle is crucial for the fixation of the cooperators’ homozygote population, as it can never occur in the case of alternating prisoner’s dilemma (see Tables 6, 9 and 12), regardless of the inheritance system. However, in the case of collaborating prisoner’s dilemma, where cooperation maximizes the sum of siblings’ survival rates, the fixation of the cooperator genotype is possible (e.g. *A*_3_ in Table 14) but not necessary, as bistability (e.g. *A*_1_ in Table 14) and coexistence (e.g. *A*_8_ in Table 14), moreover, the global stability of homozygous defector genotype (e.g. *A*_6_ in Table 14) can also occur. Therefore, it can be said that although the principle of “family welfare” is important, the fixation of the cooperator genotype does not solely depend on it.

It is well-known that theories based on group selection often rely on at least two ingredients for the spread of cooperation [30]: a) the synergetic effect of the number of cooperators on the individual fitness of group members, and b) the formation process of groups and their composition. In the case of familial selection discussed in this article, the family constitutes the group, and its composition is determined by the genetic system, but there is no presence of synergetic effects, highlighting that the process of group formation alone may be sufficient for the fixation of the cooperative phenotype.

We note that standard models of group selection focus primarily on asexual populations (e.g. [31], thus they are generally not suitable, or only limitedly applicable, for analyzing effects that arise from group formation based on genetic foundations.

### Our results from the viewpoint of kin selection

Donation game establishes a connection between the prisoner’s dilemma and the classical Hamilton’s rule (e.g. [25-27]). It belongs to the category of collaborative prisoner’s dilemma games and involves only two parameters: cost and benefit. It is found that regardless of the genotype-phenotype mapping, the genotype dynamics display the fixation of a population consisting solely of pure cooperators when the classical Hamilton’s rule is satisfied.

However, when considering prisoner’s dilemma games that typically depend on more than two parameters, a population consisting exclusively of homozygous defectors can also become fixed (see *A*_6_ in Table 14), but bistability and coexistence can occur as well. Moreover, even a change in the genotype-phenotype mapping alone can lead to a change in the stability of the population of cooperator homozygotes (see *A*_4_ in Table 14), despite the payoff matrix remaining unchanged and relatedness being 1/2. It is worth noting that relatedness is not always a necessary condition for the stability of a population consisting solely of cooperator homozygotes, as seen in the case of inequalities (12) and (14) where there is no multiplicative factor of 2, though, in a monogamous population, the relatedness among siblings is 1/2.

From the perspective of inclusive fitness, the cooperative gene can enhance its evolutionary success by indirectly contributing to the survival of siblings who have a 1/2 probability of carrying the same cooperative gene. This indirect effect also operates in the case of familial selection. In models of kin selection, the consideration of inclusive fitness explains the spread of the cooperative trait. Following this pattern, the question arose as to whether there is a mechanism or condition within genotype dynamics that guarantees an increase in the frequency of the cooperative allele along any trajectory. This led to the introduction of Haldane monotonicity. However, in contrast to the inclusive fitness approach, genotype dynamics are not only suitable for examining when a homozygote type becomes fixed, but also for addressing bistability and the coexistence of different genotypes.

In summary, it can be stated that some of our findings are consistent with the results of detailed models of kin selection and group selection (e.g., the case of the donation game). However, there are also differences, among which one of the most important is that our model highlights the significant influence of genotype-phenotype mapping on the outcome of natural selection. This may be attributed to an essential feature of our model, which takes into account diploidy, in contrast to the detailed models of kin selection, which, as noted on p. 414 of Van Veelen et al. [27], are mostly haploid models (see e.g. [29]).

### Genotype dynamics brings together the basic ideas of the kin and the group selection theories for familial selection

As mentioned above, the “welfare of the family”, i.e., the sum of the survival rates of siblings, is a crucial determinant of selection outcome. For instance, in the case of alternating prisoner’s dilemma games, the homozygous genotype of the cooperator can never fixate (as the total payoff is higher when one of the interacting siblings is a defector). On the other hand, solely based on the metric of “family welfare”, the fixation of a purely cooperative population cannot be predicted, as illustrated by the *A*_6_ payoff matrix in Table 14, where regardless of the genotype-phenotype mapping, the population always fixates to a purely defector state. Regarding the donation game, we have demonstrated that the classical Hamilton’s rule predicts the fixation of a population consisting only of pure cooperators. However, an important question arises: what is the significance of the multiplier 2 in the inequality *b* > 2*c* ([32-33]? According to the classical Hamilton’s rule, it arises from genetic relatedness. From the perspective of group selection, it represents the number of players in the matrix game (see Section 4). Both interpretations appear reasonable in the context of the donation game.

In a diploid population, if two interacting siblings aim at maximizing the “welfare of the family”, both direct and indirect effects come into play. In other words, in the case of familial selection, both kin selection and group selection operate simultaneously. Therefore, it can be stated that the current familial selection model, which focuses only on changes in genotype frequencies, reconciles the fundamental ideas of both theories under the umbrella of the “orthodox” Darwinian view.

### Conclusion

The genotype dynamics directly tracks changes in genotype distribution over time. It is not based on either the Price equation [34], nor the concept of individual-based fitness or inclusive fitness [35], nor the welfare of families [31]. The genotype dynamics can handle the Darwinian struggle for life, where the reproductive success of parents and the survival of offspring jointly determine the success of each genotype. Although this dynamics is an extension of the well-known replicator dynamics, it falls within the realm of classical population genetic models. The specific details of the genetic system determine the genotypic composition within each family, while the genotype-phenotype mapping determines the phenotypic composition.

One of our main observations is that the mathematical methods developed for classical asexual matrix games [21] can be successfully applied to population genetic models. In particular, the static definition of ESGD implies the local stability of the genotype dynamics (cf. [36-37], Theorem 7.2.4 in [21]). Furthermore, the genotype dynamics implicitly includes genetic systems where the genotype-phenotype mapping is well-defined, and the genotypes of parents unambiguously determine the genotypes of offspring. However, due to the large number of possible genotypes, applying this approach often involves high-dimensional nonlinear differential equations. We hope that Theorem 1 presented here, along with the static definition for evolutionary stability and Haldane monotonicity, will aid in both analytical and numerical investigations of other selection regimes (see Remarks 1, 2 and 3) in future studies.

During the preparation of the applications, we started from Haldane’s familial selection scenario [1]. We sought to answer the question of which phenotype can survive within a given monogamous family, assuming that the survival rates of siblings are determined by a game between them. Our approach provides the most direct way to determine which behaviors will be selected among siblings as a consequence of the game, treating the population genetic components of the model with mathematical rigor, following the advice of Van Veelen et al. [38] by considering detailed conditions. We examined the simplest genetic system: a two-allele autosomal locus with Mendelian dominant-recessive or intermediate inheritance. Assuming an arbitrary payoff matrix, we determined the conditions under which a population consisting only of homozygotes is stable and when coexistence of individuals with different genotypes is possible.

As an application of our results, we looked at the case where the game between siblings is a prisoner’s dilemma. In this case, we provided concrete examples (see Tables 14 and 15) illustrating the conditions that enable the coexistence of cooperators and defectors.

Our population genetic model, as shown by the matrix examples *A*2, *A*4, *A*5, *A*7, and *A*10 in Tables 14 and 15, highlights that the genotype-phenotype mapping significantly influences the outcome of familial selection [3, 14, 18]. This is particularly interesting because, to our knowledge, neither the kin selection nor the multilevel selection models have previously demonstrated this effect of genotype-phenotype mapping.

In the overwhelming majority of eusocial animals, such as the naked mole-rat, the family means the group. Similarly, in human societies, the family serves as the basic, traditional social unit [39]. Given this, we believe that familial selection is an important framework for understanding the evolutionary origins of human altruism and cooperation. For instance, the altruism plays a significant role in family firms in the United States (e.g. [40]). Bergstrom [41] highlighted that the documented examples of familial altruism in human families may have evolutionary roots.

Finally, from the perspective of kin selection and group selection, our monogamous population genetic model on familial selection can be considered as a special case study, in which the group is formed by the genetically well-defined family and the survival of siblings is exclusively influenced by the interactions between them. However, we note that our model is one of the simplest biological models for kin selection in diploid, sexual populations. Consequently, any comprehensive kin selection theory should include our model as a specific case.

## Materials and Methods

We introduced the general genotype dynamics (1) and notion of evolutionarily stability for the diploid genotypes. Following the standard reasoning of classical game theory, we proved that condition of the evolutionary stable genotype distribution implies the local stability of genotype dynamics. Consequently, the static condition of evolutionary stability helps in our dynamical studies. Throughout our investigation, we employ standard mathematical analysis, Lyapunov method. The figures were created using Wolfram Mathematica 12. The orbits seen in the phase portraits are the results of numerical solutions (NDSolve).

## Author Contributions

Conceptualization: JG. Methodology: JG, TFM. Formal analysis: VCs, TFM, TV. Visualization and numerical investigation: ASz, TV. Writing – Review & Editing: JG, ASz TV, VCs, TFM

## References

1. Haldane JBS. A mathematical theory of natural and artificial selection-I. Trans Camb Philos Soc. 1924;23: 19–41.

2. Theodorou K, Couvet D. Familial versus mass selection in small populations. Genet Sel Evol. 2003;35: 425. doi: 10.1186/1297-9686-35-5-425.

3. Garay J, Garay BM, Varga Z, Csiszár V, Móri TF. To save or not to save your family member’s life? Evolutionary stability of self-sacrificing life history strategy in monogamous sexual populations. BMC Evol Biol. 2019;19: 147. doi: 10.1186/s12862-019-1478-0.

4. Garay, J, Csiszár V, Móri TF. Subsistence of sib altruism in different mating systems and Haldane’s arithmetic. J Theor Biol. 2023;557: 111330. doi: 10.1016/j.jtbi.2022.111330.

5. Fisher RA. The genetical theory of natural selection. 2nd ed. Dover, New York: Oxford University Press; 1958. ISBN: 9780198504405.

6. Maynard Smith J. Evolutionary genetics. 2nd ed. Oxford University Press; 1989. ISBN-13. 978-0198502319.

7. Nagylaki T. Introduction to theoretical population genetics. Berlin: Springer-Verlag; 1994. ISBN-13. 978–0387533445.

8. Cavalli-Sforza LL, Feldman MW. Darwinian selection and “altruism”. Theor Popul Biol. 1978;14: 268–280. doi: 10.1016/0040-5809(78)90028-X.

9. Haldane JBS, Jayakar SD. Selection for a single pair of allelomorphs with complete replacement. J Genet. 1965;59: 81–87. doi: 10.1007/BF02984147.

10. Hull P. Partial incompatibility not affecting total litter size in the mouse. Genetics. 1964;50: 563–570. doi: 10.1093/genetics/50.4.563.

11. King JL. The effect of litter culling – or family planning – on the rate of natural selection. Genetics. 1965;51: 425–429. doi: 10.1093/genetics/51.3.425.

12. Maynard Smith J. Models of the evolution of altruism. Theor Popul Biol. 1980;18: 151–159. doi: 10.1016/0040-5809(80)90046-5.

13. Scudo FM, Ghiselin MT. Familial selection and the evolution of social behavior. J Genet. 1975;62: 1–31. doi: 10.1007/BF02984178.

14. Uyenoyama M, Feldman MW. Theories of kin and group selection: A population genetics perspective. Theor Popul Biol. 1980;17: 380–414. doi: 10.1016/0040-5809(80)90033-7.

15. Uyenoyama M, Feldman MW, Mueller LD. Population genetic theory of kin selection: Multiple alleles at one locus. Proc Natl Acad Sci U S A. 1981;78: 5036–5040. doi: 10.1073/pnas.78.8.5036.

16. Hamilton WD. The genetical evolution of social behaviour. I. J Theor Biol. 1964;7: 1–16. doi: 10.1016/0022-5193(64)90038-4.

17. Bergstrom TC. On the Evolution of Altruistic Ethical Rules for Siblings. Am Econ Rev. 1995;85: 58–81.

18. Toro M, Abugov R, Charlesworth B, Michod RE. Exact versus heuristic models of kin selection. J Theor Biol. 1982;97: 699–713. doi: 10.1016/0022-5193(82)90368-X.

19. Poundstone W. Prisoner’s Dilemma: John von Neumann, Game Theory, and the Puzzle of the Bomb. 1st ed. New York: Anchor; 1993. ISBN 0-385-41580-X.

20. Hilbe C, Nowak MA, Sigmund K. Evolution of extortion in Iterated Prisoner’s Dilemma games. Proc Natl Acad Sci U S A. 2013;110: 6913–8. doi: 10.1073/pnas.1214834110.

21. Hofbauer J, Sigmund K. Evolutionary games and population dynamics. Cambridge: Cambridge University Press; 1998. ISBN-13: 978-0521625708.

22. Alós-Ferrer C, Hofbauer J. Excess payoff dynamics in games. J Econ Theory. 2022;204: 105464. doi: 10.1016/j.jet.2022.105464.

23. Maynard Smith J, Price GR. The logic of animal conflict. Nature. 1973;246: 15–18. doi: 10.1038/246015a0.

24. Garay J, Varga Z. Coincidence of ESAD and ESS in dominant-recessive hereditary systems. J Theor Biol. 2003;222: 297–305. doi: 10.1016/S0022-5193(03)00027-4.

25. Doebeli M, Hauert C. Models of cooperation based on the Prisoner’s dilemma and snowdrift games. Ecol Lett. 2005;8: 748–766. doi: 10.1111/j.1461-0248.2005.00773.x.

26. Van Veelen M. The replicator dynamics with n players and population structure. J Theor Biol. 2011;276: 78–85. doi: 10.1016/j.jtbi.2011.01.044.

27. Van Veelen M, Allen B, Hoffman M, Simon B, Veller C. Hamilton’s rule. J Theor Biol. 2017;414: 176–230. doi: 10.1016/j.jtbi.2016.08.019.

28. Price GR. Selection and covariance. Nature. 1970;227: 520–521. doi: 10.1038/227520a0.

29. Gardner A, West SA, Wild G. The genetical theory of kin selection. J Evol Biol. 2011;24: 1020–1043. doi: 10.1111/j.1420-9101.2011.02236.x.

30. Simon B. Continuous-time models of group selection, and the dynamical insufficiency of kin selection models. J Theor Biol. 2014;349:22–31.

31. Nowak MA, Tarnita CE, Wilson EO. The evolution of eusociality. Nature. 2010;466: 1057–1062. doi: 10.1038/nature09205.

32. Van Veelen M. The group selection-inclusive fitness equivalence claim: Not true and not relevant. Evol Hum Sci. 2020;2: e11. doi: 10.1017/ehs.2020.9.

33. Van Veelen M, García J, Sabelis MW, Egas M. Group selection and inclusive fitness are not equivalent; the Price equation vs. models and statistics. J Theor Biol. 2012;299: 64–80. doi: 10.1016/j.jtbi.2011.07.025.

34. Van Veelen M. On the use of the Price equation. J Theor Biol. 2005;237: 412–426. doi: 10.1016/j.jtbi.2005.04.026.

35. Abbot P, Abe J, Alcock J, et al. Inclusive fitness theory and eusociality. Nature. 2011;471: E1–E4. doi: 10.1038/nature09831.

36. Hofbauer J, Schuster P, Sigmund K. A note on evolutionarily stable strategies and game dynamics. J Theor Biol. 1979;81: 609–612. doi: 10.1016/0022-5193(79)90058-4.

37. Zeeman EC. Population dynamics from game theory. In: Nitecki Z, Robinson C, editors. Global Theory of Dynamical Systems. Lecture Notes in Mathematics, vol 819. Berlin, Heidelberg: Springer; 1980. pp. 471–497. doi: 10.1007/BFb0087009.

38. Van Veelen M, García J, Sabelis MW, Egas M. Call for a return to rigour in models. Nature. 2010;467: 661. doi: 10.1038/467661d.

39. Geary DC, Flinn MV. Evolution of Human Parental Behavior and the Human Family. Parent Sci Pract. 2001;1: 5–61. doi: 10.1080/15295192.2001.9681209.

40. Schulze WS, Lubatkin MH, Dino RN. Toward a theory of agency and altruism in family firms. J Bus Ventur. 2003;18:473–90. doi: 10.1016/S0883-9026(03)00054-5.

41. Bergstrom TC. Economics in a Family Way. J Econ Lit. 1996;34: 1903–34.

